# RPW8/HR Repeats Control NLR Activation in *A. thaliana*

**DOI:** 10.1101/559864

**Authors:** Cristina A. Barragan, Rui Wu, Sang-Tae Kim, Wanyan Xi, Anette Habring, Jörg Hagmann, Anna-Lena Van de Weyer, Maricris Zaidem, William Wing Ho Ho, George Wang, Ilja Bezrukov, Detlef Weigel, Eunyoung Chae

## Abstract

In many plant species, conflicts between divergent elements of the immune system, especially nucleotide-binding oligomerization domain-like receptors (NLR), can lead to hybrid necrosis. Here, we report deleterious allele-specific interactions between an NLR and a non-NLR gene cluster, resulting in not one, but multiple hybrid necrosis cases in *Arabidopsis thaliana*. The NLR cluster is *RESISTANCE TO PERONOSPORA PARASITICA 7* (*RPP7*), which can confer strain-specific resistance to oomycetes. The non-NLR cluster is *RESISTANCE TO POWDERY MILDEW 8* (*RPW8*) / *HOMOLOG OF RPW8* (*HR*), which can confer broad-spectrum resistance to both fungi and oomycetes. RPW8/HR proteins contain at the N-terminus a potential transmembrane domain, followed by a specific coiled-coil (CC) domain that is similar to a domain found in pore-forming toxins MLKL and HET-S from mammals and fungi. C-terminal to the CC domain is a variable number of 21- or 14-amino acid repeats, reminiscent of regulatory 21-amino acid repeats in fungal HET-S. The number of repeats in different RPW8/HR proteins along with the sequence of a short C-terminal tail predicts their ability to activate immunity in combination with specific RPP7 partners. Whether a larger or smaller number of repeats is more dangerous depends on the specific RPW8/HR autoimmune risk variant.

**Author Summary:** In many plant species, conflicts between divergent elements of the immune system can cause hybrids to express autoimmunity, a generally deleterious syndrome known as hybrid necrosis. We are investigating multiple hybrid necrosis cases in *Arabidopsis thaliana* that are caused by allele-specific interactions between different variants at two unlinked resistance (R) gene clusters, *RESISTANCE TO PERONOSPORA PARASITICA 7* (*RPP7*) and *RESISTANCE TO POWDERY MILDEW 8* (*RPW8*)/*HOMOLOG OF RPW8* (*HR*). The *RPP7* locus encodes intracellular nucleotide binding site-leucine rich repeat (NLR) immune receptors that can confer strain-specific resistance to oomycetes, while the *RPW8*/*HR* locus encodes atypical resistance proteins, of which some can confer broad-spectrum resistance to filamentous pathogens. There is extensive structural variation in the *RPW8/HR* cluster, both at the level of gene copy number and at the level of C-terminal, 21- or 14-amino acid long RPW8/HR repeats. We demonstrate that the number of RPW8/HR repeats and the short C-terminal tail correlate, in an allele-specific manner, with the severity of hybrid necrosis when these alleles are combined with *RPP7* variants. We discuss these findings in light of sequence similarity between RPW8/HR and pore-forming toxins MLKL and HET-S from mammals and fungi.

## Introduction

The combination of divergent parental genomes in hybrids can produce new phenotypes not seen in either parent. At one end of the spectrum is hybrid vigor, with progeny being superior to the parents, while at the other end there is hybrid weakness, with progeny being inferior to the parents, and in the most extreme cases being sterile or unable to survive.

In plants, a particularly conspicuous set of hybrid incompatibilities is associated with autoimmunity, often with substantial negative effects on hybrid fitness [1–3]. Studies of hybrid autoimmunity in several species, often expressed as hybrid necrosis, have revealed that the underlying genetics tends to be simple, with often only one or two major-effect loci. Where known, at least one of the causal loci encodes an immune protein, often an intracellular nucleotide binding site-leucine-rich repeat (NLR) protein [4–13]. The gene family encoding NLR immune receptors is the most variable gene family in plants, both in terms of inter- and intraspecific variation [14–17]. Many NLR proteins function as major disease resistance (R) proteins, with the extravagant variation at these loci being due to a combination of maintenance of very old alleles by long-term balancing selection and rapid evolution driven by strong diversifying selection [18–20]. The emergence of new variants is favored by many NLR genes being organized in tandem clusters, which can spawn new alleles as well as copy number variation by illegitimate recombination, and by the presence of leucine rich repeats in NLR genes, which can lead to expansion and contraction of coding sequences [21–23]. Cluster expansion has been linked to diversification and adaptation in a range of systems [24–26]. Several complex plant NLR loci provide excellent examples of cluster rearrangement increasing pathogen recognition specificities [19]. Substantial efforts have been devoted to decomposing the complexity of the plant immune system and interactions between its components.

While many plant disease R genes are members of the NLR family, some feature different molecular architectures. One of these is *RESISTANCE TO POWDERY MILDEW 8 (RPW8)* in *Arabidopsis thaliana*, which was first identified based on an allele that confers resistance to multiple powdery mildew isolates [27]. The namesake *RPW8* gene is located in a gene cluster of variable size and composition that includes multiple RPW8-like genes as well as *HOMOLOG OF RPW8 (HR*) genes [27–29]. The reference accession Col-0, which is susceptible to powdery mildew, has four *HR* genes, but no *RPW8* gene, whereas the resistant accession Ms-0 carries *RPW8.1* and *RPW8.2* along with three *HR* genes [27]. NLRs are distinguished by N-terminal Toll/interleukin-1 receptor (TIR) or coiled-coil (CC) domains, which, when overexpressed alone, can often activate immune signaling [30,31]. A subset of CC-NLRs (CNLs) has a diagnostic type of coiled-coil domain, termed CC_R_ to indicate that this domain is being shared with RPW8/HR proteins. The latter have an N-terminal extension that might be a transmembrane domain as well as C-terminal repeats of unknown activity [32,33]. It has been noted that the CC_R_ domain is similar to a portion of the animal mixed-lineage kinase domain-like (MLKL) protein that forms a multi-helix bundle [34] as well as the HeLo and HELL domains of fungi, which also form multi-helix bundles [35–37]. Many fungal HeLo domain proteins have a prion-forming domain that consists of C-terminal 21-amino acid repeats. This domain can form amyloids and thereby affect oligomerization and activity of these proteins [35–39].

We have previously reported hybrid necrosis due to incompatible alleles at the *RPW8/HR* locus and at the complex *RECOGNITION OF PERONOSPORA PARASITICA 7 (RPP7)* locus, which encodes a canonical CNL and which has alleles that provide race-specific resistance to the oomycete *Hyaloperonospora arabidopsidis* [54,55]. Here, we investigate in detail three independent cases of incompatible *RPW8/HR* and *RPP7* alleles, and show that two are caused by members of the fast-evolving RPW8.1/HR4 clade. We describe how variation in the number of C-terminal repeats and the short C-terminal tail predict the degree of incompatibility between two common *RPW8.1/HR4* alleles and corresponding *RPP7* alleles.

## Results

### Distinct pairs of RPP7 and RPW8/HR alleles cause hybrid necrosis

In a systematic intercrossing and genetic mapping program among 80 *A. thaliana* accessions, a series of genomic regions involved in hybrid incompatibility were identified. The underlying genes were termed *DANGEROUS MIX* (*DM*) loci. One instance, between the *DM6* and *DM7* regions, stood out because it is responsible for two phenotypically distinct hybrid necrosis cases (**Fig 1A**) [10]. Strong candidates, as previously inferred from a combination of mapping, gene knockdown and transformation with genomic constructs, suggested that *DM6* corresponds to the *RPP7* cluster, and *DM7* to the *RPW8*/*HR* cluster. We recently found an additional case of incompatibility between the *DM6* and *DM7* regions, with a third distinctive phenotype (**Fig 1A and 2A**). In addition to phenotypic differences between the three *DM6*–*DM7* F_1_ hybrids, test crosses confirmed that each case was caused by different combinations of *DM6* and *DM7* alleles, as only certain combinations resulted in hybrid necrosis (**Fig 1B**).

**Fig 1.**
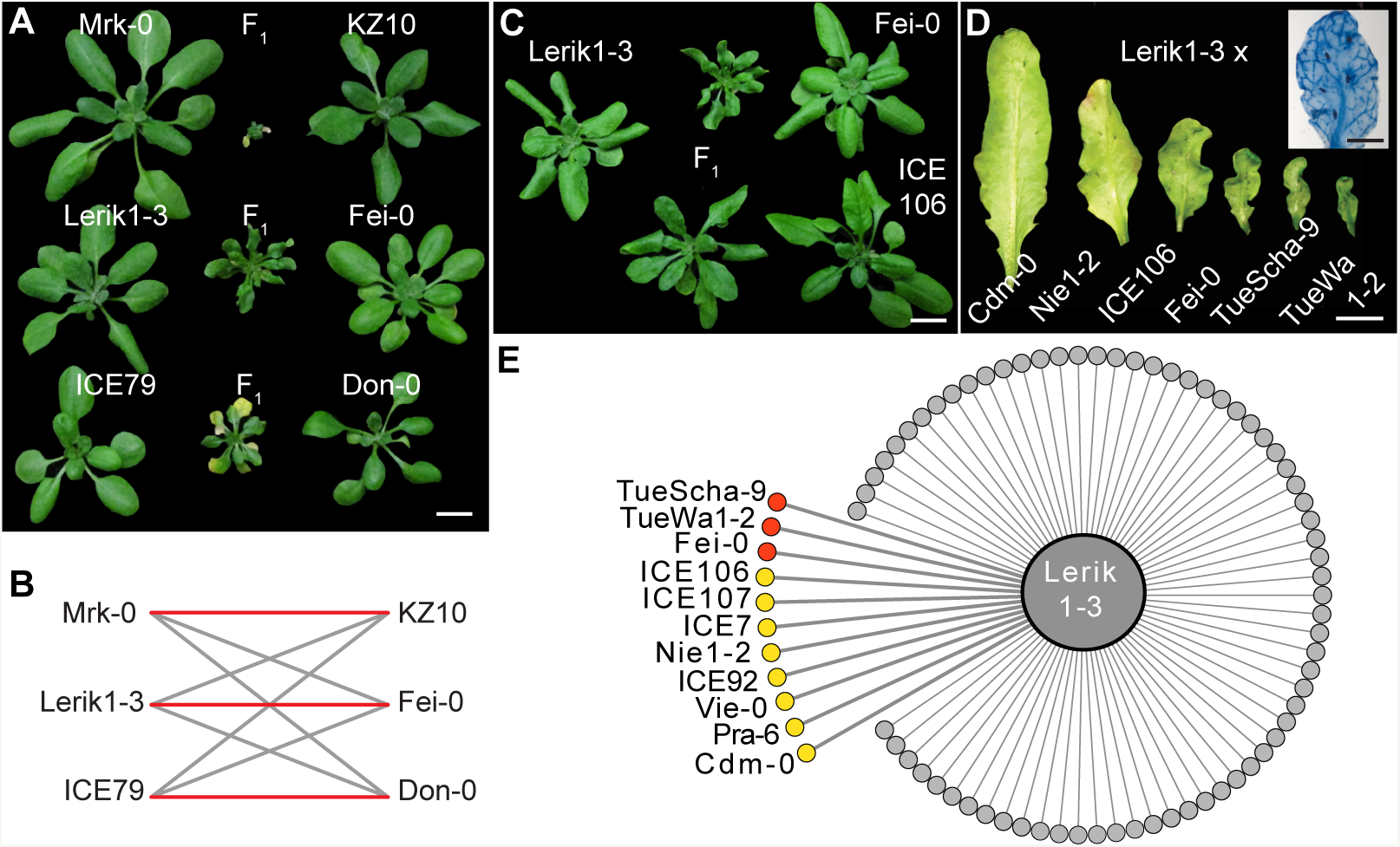
*DM6*–*DM7* hybrid necrosis cases. **(A)** Morphological variation in three independent *DM6*–*DM7* hybrid necrosis cases. **(B)** Red lines indicate necrosis in F_1_ hybrids, black indicates normal progeny. **(C, D)** Variation in morphology in two *DM6*–*DM7* cases sharing the same *DM6* allele in Lerik1-3. **(C)** Entire rosettes of four-week-old plants. **(D)** Abaxial sides of eighth leaves of six-week-old plants. Inset shows Trypan Blue stained leaf of Lerik1-3 x Fei-0 F_1_. **(E)** Summary phenotypes in crosses of Lerik1-3 to 80 other accessions. Red is strong necrosis in F_1_, and yellow is mild necrosis in F_1_ or necrosis only observable in F_2_. Scale bars indicate 1 cm.

To corroborate the evidence from mapping experiments that *DM6* alleles of Mrk-0 and ICE79 were *RPP7* homologs, we designed ten artificial microRNAs (amiRNAs) based on sequences from the Col-0 reference accession. AmiRNAs targeting a subclade of five *RPP7* homologs that make up the second half of the *RPP7* cluster in Col-0, suppressed hybrid necrosis in all three crosses, Mrk-0 x KZ10, Lerik1-3 x Fei-0 and ICE79 x Don-0 (**Fig S1** and **Table S1**). These rescue experiments, together with the above-mentioned test crosses, indicate that specific *RPP7* homologs in Mrk-0, Lerik1-3 and ICE79 correspond to different *DM6* alleles that cause hybrid necrosis in combination with specific *DM7* alleles from other accessions.

### A common set of *RPW8*/*HR* haplotypes affecting hybrid performances in F_1_ and F_2_ progeny

In the mentioned set of diallelic F_1_ crosses among 80 accessions [10], we noted that the *DM6* carrier Lerik1-3 was incompatible with several other accessions, suggesting that these have *DM7* (*RPW8*/*HR*) hybrid necrosis risk alleles that are similar to the one in Fei-0. Crosses with TueScha-9 and TueWa1-2 produced hybrids that looked very similar to Lerik1-3 x Fei-0 progeny, with localized spots of cell death spread across the leaf lamina along with leaf crinkling and dwarfism (**Fig 1D and S2**). Similar spots of cell death and leaf crinkling were observed in crosses of Lerik1-3 to ICE106 and ICE107, although these were not as dwarfed (**Fig 1C,D and S2**).

Hybrid necrosis often becomes more severe when the causal loci are homozygous [5,7,10,12]. To explore whether Lerik1-3 might cause milder forms of hybrid necrosis that are missed in the F_1_ generation, we surveyed several F_2_ populations involving Lerik1-3. Six segregated necrotic plants with very similar phenotypes (**Fig 1D,E** and **S2**). This makes all together for 11 incompatible accessions, which are spread over much of Eurasia (**Fig 1E**).

The F_2_ segregation ratios suggested that the *DM7* allele from ICE106/ICE107 is intermediate between the Fei-0/TueWa1-2/TueScha-9 alleles and the Cdm-0/Nie-0 alleles **(Table 1)**. Alternatively, the hybrid phenotypes might be affected by background modifiers, such that identical *DM7* alleles produce a different range of phenotypes in combination with *DM6*^Lerik1-3^.

**Table 1:**
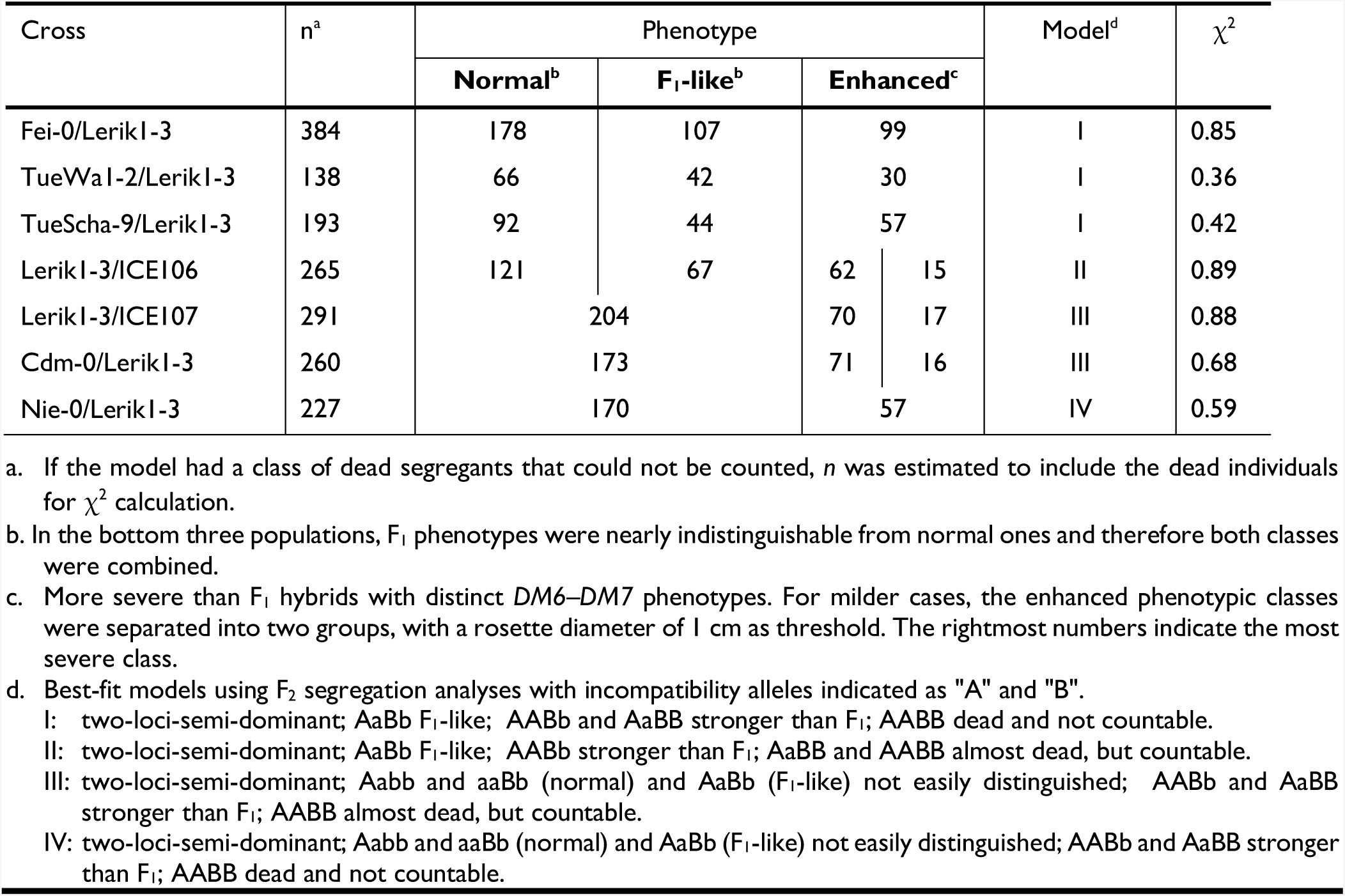
F_2_ segregation ratios at 16 °C.

Because the phenotypic variation among hybrid necrosis cases involving Lerik1-3 could involve loci other than *DM6* and *DM7*, we carried out linkage mapping with Lerik1-3 x ICE106 and Lerik1-3 x ICE107 crosses. We combined genotyping information from Lerik1-3 x ICE106 and Lerik1-3 x ICE107 F_2_ and F_3_ individuals for mapping, because the genomes of ICE106 and ICE107, which come from closeby collection sites, are very similar and because the two crosses produce very similar F_1_ hybrid phenotypes, suggesting that the responsible alleles are likely to be identical. We used F_3_ populations to better distinguish different phenotypic classes, since we did not know the number of causal genes nor there genetic behavior. QTL analysis confirmed that the *DM6* and *DM7* genomic regions are linked to hybrid necrosis in these crosses (**Fig 2A,B**).

**Fig 2.**
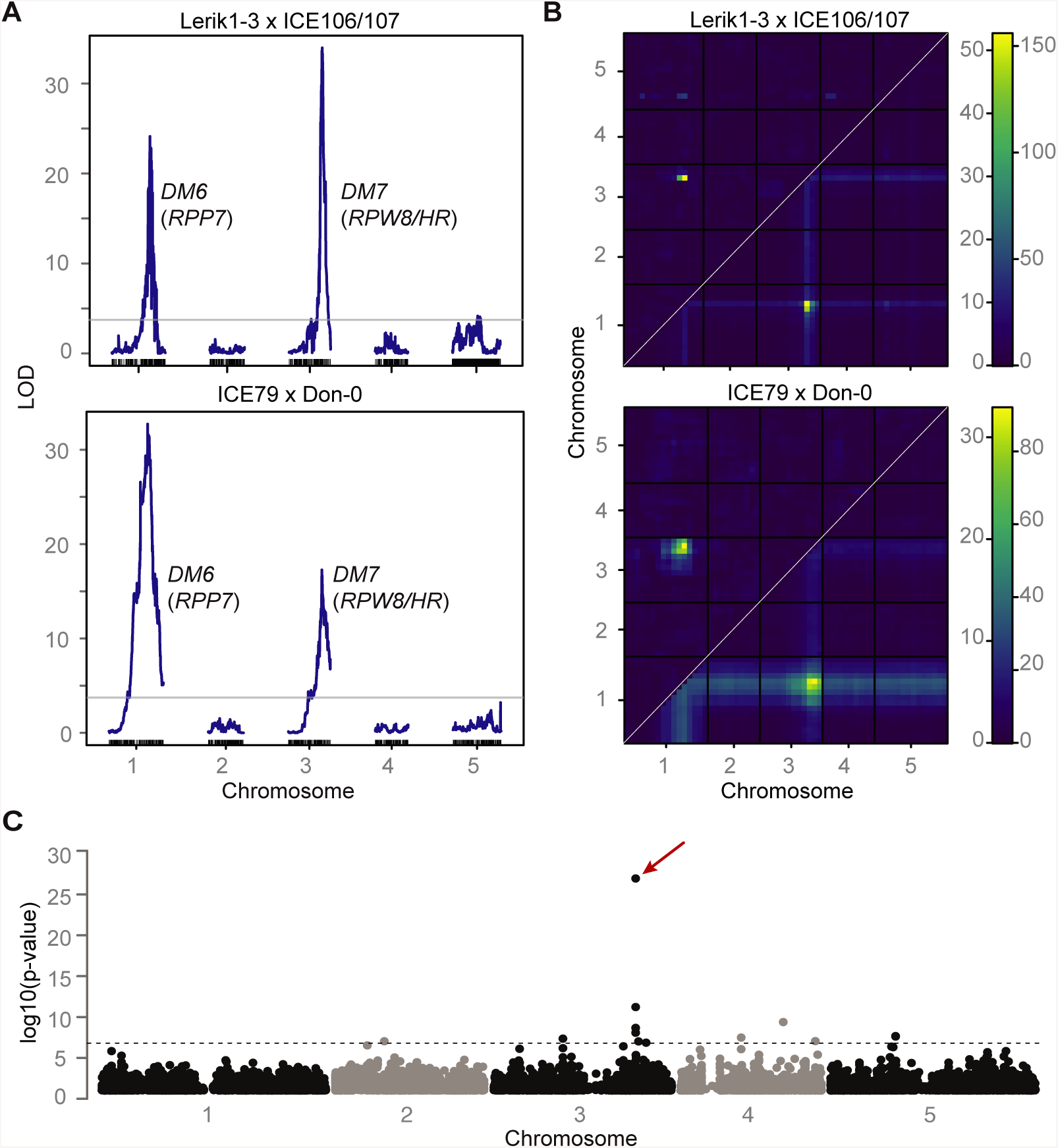
Mapping of two *DM6*–*DM7* hybrid necrosis cases. **(A)** QTL analyses. The QTL on chromosome 1 includes *RPP7* from Lerik1-3 and ICE79 (21.37-22.07 and 21.50-21.98 Mb), and the QTL on chromosome 3 *RPW8*/*HR* from ICE106/ICE107 and Don-0 (18.59-19.09 Mb, 18.61-19.06 Mb). The horizontal lines indicate 0.05 significance thresholds established after 1,000 permutations. **(B)** Heat map for two-dimensional, two-QTL model genome scans. Upper left triangles indicate epistasis scores (LOD*i*) and lower right triangles joint two-locus scores (LOD*f*). Scales for LOD*i* on left and for LOD*f* on right. **(C)** Manhattan plot for a GWAS of necrosis in hybrid progeny of Lerik1-3 crossed to 80 other accessions (see Table S2). The hit in the *RPW8*/*HR* region (red arrow) stands out, but It is possible that some of the other hits that pass the significance threshold (Bonferroni correction, 5% familywise error) identify modifiers of the *DM6*–*DM7* interaction.

To narrow down the *DM7* mapping interval, we took advantage of having 11 accessions that produced hybrid necrosis in combination with Lerik1-3, and 69 accessions (including Lerik1-3 itself) that did not. We performed GWAS with Lerik1-3-dependent hybrid necrosis as a binary trait [40]. The by far most strongly associated marker was immediately downstream of *HR4*, the last member of the *RPW8*/*HR* cluster in Col-0 (**Fig 2C** and **Table S2**). An amiRNA matching *HR4* sequences from Col-0 fully rescued both the strong necrosis in Lerik1-3 x Fei-0 and the weaker necrosis in Lerik1-3 x ICE106 (**Fig 3A** and **Table S3**). We confirmed the causality of another member of the *RPW8*/*HR* cluster in the KZ10 x Mrk-0 case with a CRISPR/Cas9-induced mutation of *RPW8.1*^KZ10^ (**Fig 3B** and **Fig S3**).

**Fig 3.**
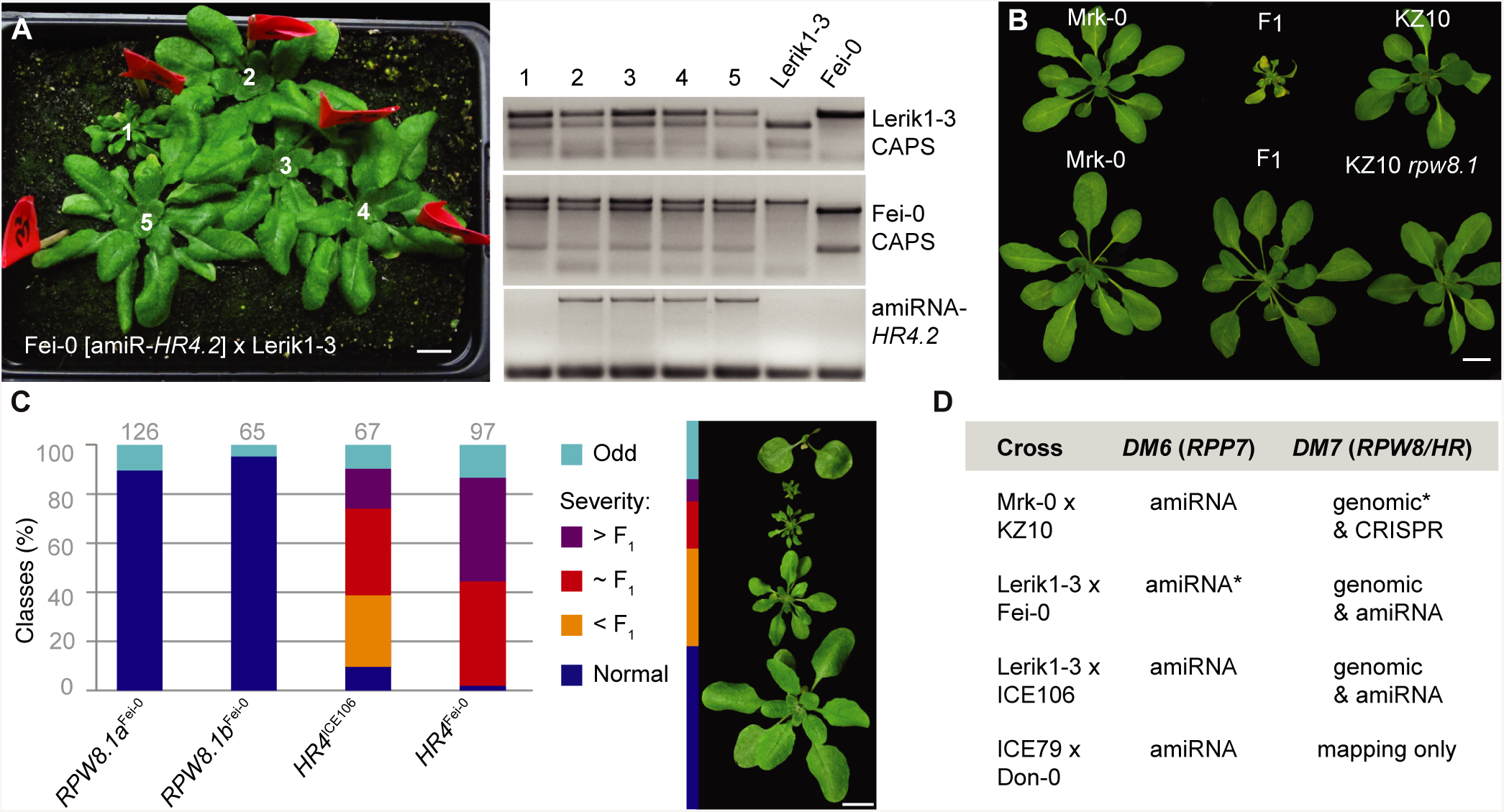
Confirmation of causal genes in *RPW8*/*HR* cluster. **(A)** Rescue of hybrid necrosis in Lerik1-3 x Fei-0 F_1_ plants with an amiRNA against *HR4*. Fei-0 parents were T_1_ transformants. PCR genotyping of numbered plants from left shown on the right. Only plant 1, which does not carry the amiRNA, is necrotic and dwarfed. **(B)** Rescue of hybrid necrosis in Mrk-0 x KZ10 F_1_ plants by CRISPR*/*Cas9-targeted mutagenesis on *RPW8.1*^KZ10^. **(C)** Recapitulation of hybrid necrosis in Lerik1-3 T_1_ plants transformed with indicated genomic fragments from Fei-0 and ICE106. Representative phenotypes on right. Numbers of T_1_ plants examined given on top. **(D)** Summary of rescue and recapitulation experiments. Asterisks refer to published experiments [10]. Scale bars indicate 1 cm.

Naturally, we wanted to learn what the relationship, if any, was between *RPP7*-dependent hybrid necrosis and the previously described function of certain *RPP7* alleles in conferring resistance to *H. arabidopsidis*. The *RPP7*^Col-0^ allele makes Col-0 resistant to the *H. arabidopsidis* isolate Hiks1 [41]. Lerik1-3, with the *RPP7*^Lerik1-3^ risk allele, is resistant to Hiks1 as well, while the Lerik1-3 incompatible accessions Fei-0 and ICE106 are not (**Fig S4** and **Table S4**). Both Lerik1-3 x Fei-0 and Lerik1-3 x ICE106 F_1_ hybrids were resistant to Hiks1, although apparently somewhat less so than the Lerik1-3 parents (**Fig S4**). We further tested whether *RPP7*-like genes from Lerik1-3 are likely to be involved in Hiks1 resistance by inoculating Hiks1 on transgenic lines carrying seven different amiRNAs against *RPP7* homologs (**Table S1**), but none of the amiRNAs reduced Hiks1 resistance. These negative results are difficult to interpret; the amiRNAs might not efficiently knock down all *RPP7* homologs in Lerik1-3 (for which the exact structure of the *RPP7* cluster is unknown), Hiks1 resistance in Lerik1-3 might require other *RPP* loci (of which there are many in the *A. thaliana* genome), or Hiks1 resistance might be complex, as found in other accessions [42]. Finally, given the interaction of *RPP7* with *RPW8*/*HR* in hybrid necrosis, we asked whether *HR4* is required for *RPP7-*mediated Hiks1 resistance in Col-0. Two independent *hr4* CRISPR/Cas9 knockout lines were generated in Col-0 (**Fig S3**), but both remained completely resistant to Hiks1 (**Fig S4** and **Table S4**), indicating that *HR4* in Col-0 is dispensable for *RPP7*-mediated resistance to Hiks1.

### Structural variation of the *RPW8*/*HR* cluster

For reasons of convenience, we assembled the *RPW8*/*HR* cluster from TueWa1-2 instead of Fei-0; accession TueWa1-2 interacted with *RPP7*^Lerik1-3^ in the same manner as Fei-0, the strong necrosis in Lerik1-3 x TueWa1-2 was rescued with the same amiRNA as in Lerik1-3 x Fei-0 (**Table S3**), and TueWa1-2 had an *HR4* allele that was identical in sequence to *HR4*^Fei-0^. We found that the *RPW8*/*HR* cluster from TueWa1-2 had at least 13 *RPW8*/*HR*-like genes, several of which were very similar to each other (**Fig 4A**). For example, there were at least four copies of *RPW8.3*-like genes with 93 to 99.8% sequence similarity, and two identical *RPW8.1* genes, named *RPW8.1a,* followed by distinct *RPW8*/*HR* copies.

**Fig 4.**
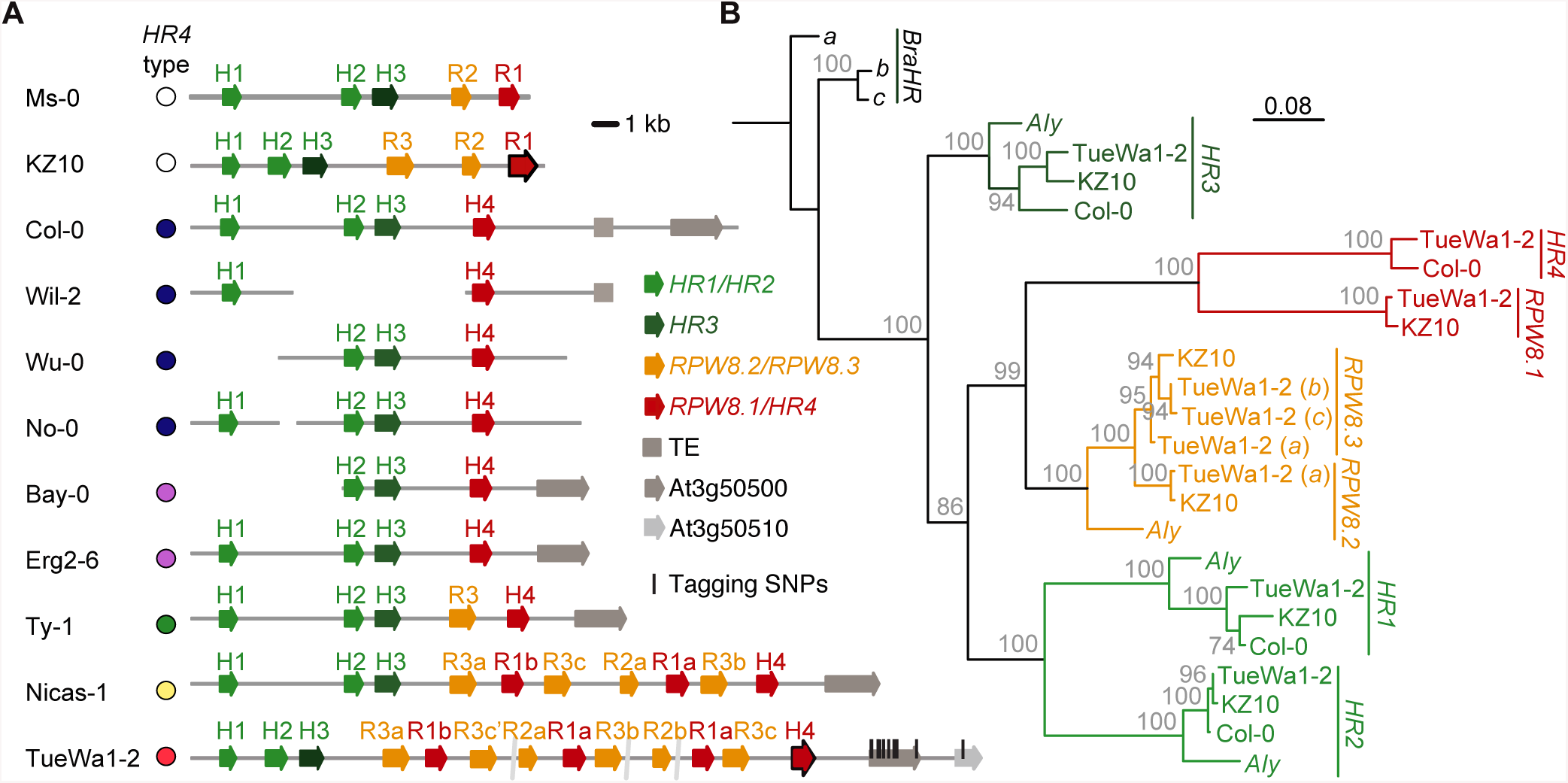
Structural variation of the *RPW8/HR* cluster. **(A)** The *RPW8/HR* cluster in different accessions. The extreme degree of recent duplications in TueWa1-2, with the same *HR4* hybrid necrosis risk allele as Fei-0, did not allow for closure of the assembly from PCR products; assembly gaps are indicated. Color coding of *HR4* alleles according to Fig 6. Tagging SNPs found in GWAS marked in TueWa1-2 *RPW8/HR* cluster as black vertical lines. **(B)** Maximum likelihood tree of *RPW8*/*HR* genes from three *A. thaliana* accessions and the *A. lyrata* and *B. rapa* reference genomes. Branch lengths in nucleotide substitutions are indicated. Bootstrap values (out of 100) are indicated on each branch.

Recapitulation experiments had identified *HR4*^Fei-0^ (identical to *HR4*^TueWa1-2^ and *HR4*^TueScha-9^) and *HR4*^ICE106^ as causal for hybrid necrosis (**Fig 3C,D**). We analyzed the phylogenetic relationship of the *RPW8*/*HR* genes in TueWa1-2 with the ones from published *RPW8*/*HR* clusters in *A. thaliana*, in *A. lyrata* and in *Brassica* spp. [10,28,29,43]. In *A. thaliana, RPW8*/*HR* genes seem to have undergone at least three duplication events, with the first one generating a new *A. thaliana* specific clade, which gave rise to independent *RPW8.1*/*HR4* and *RPW8.2*/*RPW8.3* duplications.

The *RPW8*/*HR* cluster of TueWa1-2 consists of *RPW8*/*HR* members from both the ancestral and the two *A. thaliana* specific clades, an arrangement that has not been observed before. Using species-wide data [44], we found that accessions carrying Col-0-like *HR4* alleles have simple cluster configurations, while accessions with *HR4* genes resembling hybrid necrosis alleles have more complex configurations (**Fig 4A**). The tagging SNPs found in GWAS (**Fig 4A,** and **Table S2**) were mostly found to be associated with the complex clusters, suggesting that the tagging SNPs are linked to structural variation in the distal region of the *RPW8*/*HR* cluster (**Fig 4B**).

### Causality of RPW8/HR C-terminal repeats

To further narrow down the mutations that cause autoimmunity, we compared *RPW8.1*^KZ10^ and *HR4*^Fei-0^ with other *RPW8*/*HR* alleles from the global *A. thaliana* collection [44]. Some *RPW8.1* alleles have intragenic duplications of a sequence encoding a 21-amino acid repeat (QWDDIKEIKAKISEMDTKLA[D/E]) at the C-terminal end of the protein [29]. In HR4, there is a related 14-amino acid repeat (IQV[H/D]QW[T/I]DIKEMKA). Both RPW8.1 and HR4 repeats are predicted to fold into extended alpha-helices, but only RPW8.1 repeats appear to have the potential to form coiled coils [45].

The number of repeats varies in both *RPW8.1* and *HR4* between hybrid necrosis risk and non-risk alleles. To experimentally test the effect of repeat number variation and other polymorphisms, we generated a series of derivatives in which we altered the number of repeats and swapped different portions of the coding sequences between the *RPW8.1*^KZ10^ risk and *RPW8.1*^Ms-0^ non-risk alleles, and between the *HR4*^Fei-0^ and *HR4*^ICE106^ risk and the *HR4*^Col-0^ non-risk alleles (**Fig 5A**).

**Fig 5.**
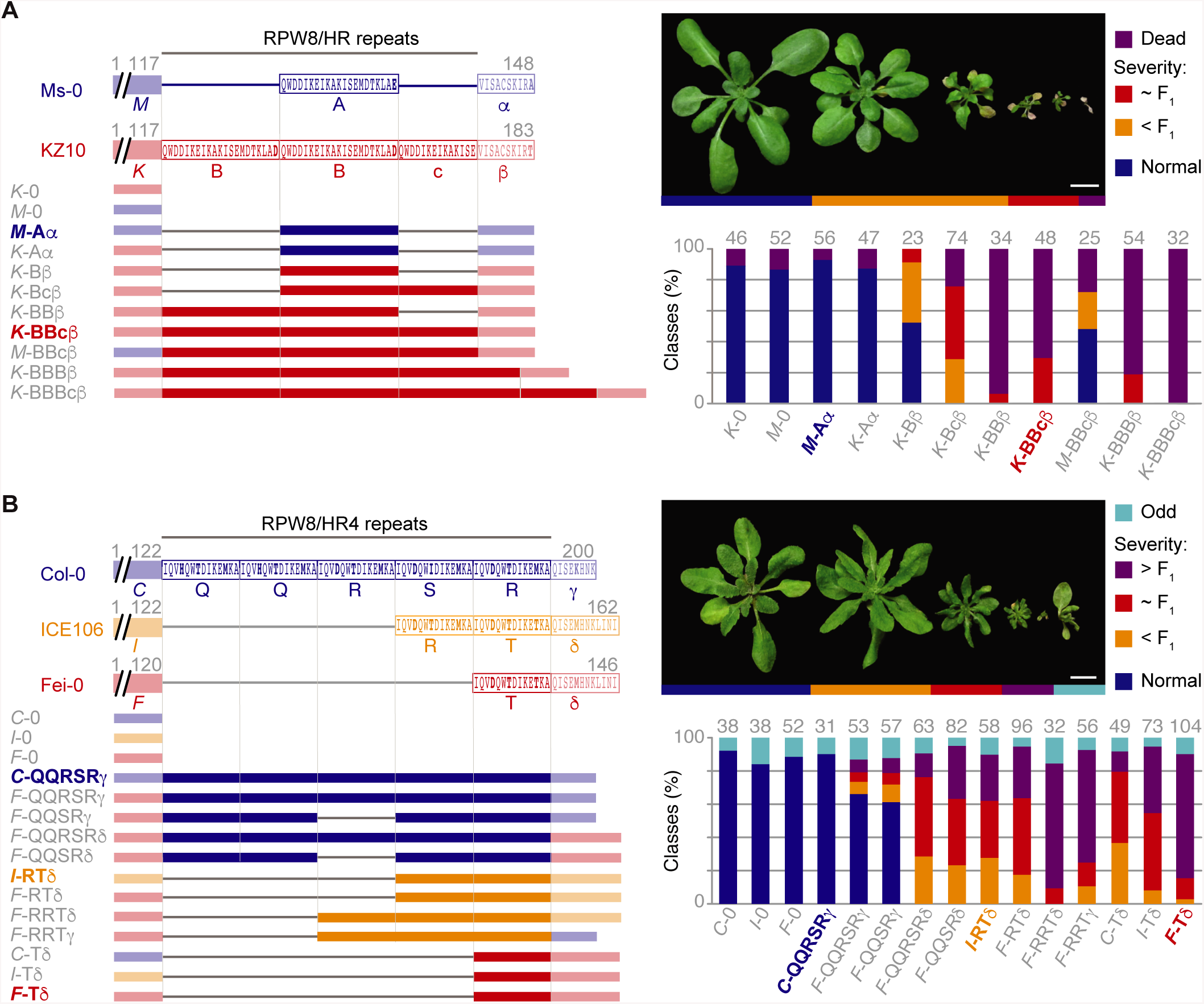
Necrosis-inducing activity of *RPW8.1* and *HR4* chimeras. N-terminal portions indicated with the initial of the accession in italics (“*K*”, “*M*”, etc.), complete repeats indicated with regular capital letters (“A”, “B”, etc.), the partial repeat in KZ10 with a lowercase letter (“c”), and the C-terminal tails with Greek letters (“α”, “β”, etc.). Non-repeat portions are semi-transparent. Repeats with identical amino acid sequences have the same letter designation. Numbers indicate amino acid positions. Constructs on the left, and distribution across phenotypic classes in T_1_ transformants on the right, with *n* given on top of each column. Natural alleles labeled in color and bold. **(A)** *RPW8.1* chimeras, driven by the *RPW8.1*^KZ10^ promoter, were introduced into Mrk-0, which carries the corresponding incompatible *RPP7* allele. **(B)** *HR4* chimeras, driven by the *HR4*^Fei-0^ promoter, were introduced into Lerik1-3, which carries the corresponding incompatible *RPP7* allele. Scale bars indicate 1 cm.

A 1.4 kb promoter fragment of *RPW8.1*^KZ10^ and a 1.2 kb promoter fragment of *HR4*^Fei-0^ in combination with coding sequences of risk alleles were sufficient to induce hybrid necrosis (**Fig 3C** and **Fig 5A, B**). To simplify discussion of the chimeras, the N-terminal portion was labeled with the initial of the accession in italics (“*M*”, “*K*”, etc.), complete repeats were labeled with different capital letters to distinguish sequence variants (“A”, “B”, etc.), the partial repeat in KZ10 with a lowercase letter (“c”), and the C-terminal tails with Greek letters (“α”, “β”, etc.).

In *RPW8.1*^KZ10^, there are two complete repeats and one partial repeat, while *RPW8.1*^Ms-0^ has only one repeat (**Fig 5A**). Modifying the number of repeats in *RPW8.1* affected the frequency and severity of necrosis in T_1_ plants in a Mrk-0 background, which carries the interacting *RPP7* allele, dramatically. Deletion of the first full repeat in *RPW8.1*^KZ10^ (“*K*-Bcβ”, with the KZ10 configuration being “*K*-BBcβ”) substantially reduced the number of plants that died in the first three weeks of growth. The additional deletion of the partial repeat (“*K*-Bβ”) reduced death and necrosis even further (**Fig 5A**). That *K*-Bβ still produces some necrosis, even though its repeat structure is the same as in the inactive *K*-Aα suggests that the polymorphism in the C-terminal tail makes some contribution to necrosis activity. It is less likely that the polymorphism in the repeats play a role, as there is only a very conservative aspartate-glutamate difference between α and β repeats.

In contrast to repeat shortening, the extension of the partial repeat (“*K*-BBBβ”) or addition of a full repeat (“*K*-BBBcβ”) increased the necrosis-inducing activity of *RPW8.1*^KZ10^, such that almost all T_1_ plants died without making any true leaves. However, it appears that not all repeats function equally, as removal of the partial repeat slightly increased necrosis-inducing activity (“*K*-BBβ”). Polymorphisms in the N-terminal non-repeat region seemed to contribute to necrosis, as swaps of the N-terminal Ms-0 fragment (“*M*-BBcβ” or “*M*-BBBβ”) induced weaker phenotypes than the corresponding variants with the N-terminal fragment from KZ10. Nevertheless, we note that the normal KZ10 repeat configuration was sufficient to impart substantial necrosis-inducing activity on a chimera in which the N-terminal half was from Ms-0, which is distinguished from KZ10 by nine nonsynonymous substitutions outside the repeats.

Compared to the RPW8.1 situation, the relationship between HR4 repeat length and necrosis-inducing activity is more complex. The natural alleles suggested a negative correlation of repeat number with necrosis-inducing activity when crossed to Lerik1-3, since the non-risk HR4 allele from Col-0 has five full repeats, while weaker risk alleles such as the one from ICE106 have two, and the strong risk allele from Fei-0 has only one (**Fig 5B**). Addition of a full repeat to HR4^Fei-0^ (“*F*-RTδ”, with the original Fei-0 configuration being “*F-*Tδ”) reduced its activity to a level similar to that of HR4^ICE106^ (“*I*-RTδ”). Deletion of a full repeat from HR4^ICE106^ (“*I*-Tδ”) modestly increased HR4 activity (**Fig 5B**). Together, the chimera analyses indicated that the quantitative differences between crosses of Fei-0 and ICE106 to Lerik1-3 (**Fig 1** and **S2**) are predominantly due to variation in HR4 repeat number. This is further supported by the necrosis-inducing activity of a chimera in which the repeats in the Col-0 non-risk allele were replaced with those from *HR4*^Fei-0^ (“*C*-Tδ”, with the original Col-0 configuration being “*C*-QQRSRγ”) (**Fig 5B** and **S5**). However, repeat number alone is not the only determinant of necrosis-inducing activity of *HR4* in combination with *RPP7*^Lerik1-3^. Adding another repeat to the “*F*-RTδ” chimera, resulting in “*F*-RRTδ”, increased the activity of *HR4*^Fei-0^ again, perhaps suggesting that there is an optimal length for *HR4* to interact with *RPP7*^Lerik1-3^.

Unlike RPW8.1, the C-terminal tails of HR4 proteins beyond the RPW8/HR repeats (fragments “γ” and “δ”) differ in length between hybrid necrosis-risk and non-risk variants (**Fig 5B**). Swapping only these two fragments affected HR4 activity substantially, and converted two chimeras with weak necrosis-inducing activity (“*F*- QQRSRγ” to “*F*-QQRSRδ” and “*F*-QQSRγ” to “*F*-QQSRδ”) into chimeras with activity resembling that of HR4^ICE106^ (which is “*I*-RTδ”).

Taken together, the swap experiments led us to conclude that naturally occurring variation in the configuration of RPW8/HR repeats play a major role in quantitatively modulating the severity of autoimmune phenotypes when these *RPW8*/*HR* variants are combined with *RPP7* alleles from Mrk-0 and Lerik1-3. At least in the case of HR4, we could show directly that the short C-terminal tail also affects the hybrid phenotype, while for RPW8.1 this seems likely as well, given that the repeats between different alleles differ less from each other than the tails.

### Prediction of *RPP7*-dependent hybrid performance using *RPW8.1*/*HR4* haplotypes

To obtain a better picture of *RPW8.1*/*HR4* variation, we remapped the raw reads from the 1001 Genomes project to the longest *RPW8.1* and *HR4* alleles, *RPW8.1*^KZ10^ and *HR4*^Col-0^, as references (**Table S5** and **S6**). The results suggested that *HR4*-carrying accessions are more rare than those carrying *RPW8.1* alleles (285 vs. 903 out of 1,221 accessions). The short, necrosis-linked, *HR4* risk alleles (**Fig 6A**) were predicted to be as frequent as the long non-risk variants (**Fig 6A, B** and **Table S5**), whereas for *RPW8.1*, only seven accessions were predicted to have the long *RPW8.1*^KZ10^-type risk variant (**Fig 6A** and **Table S6**).

**Fig 6.**
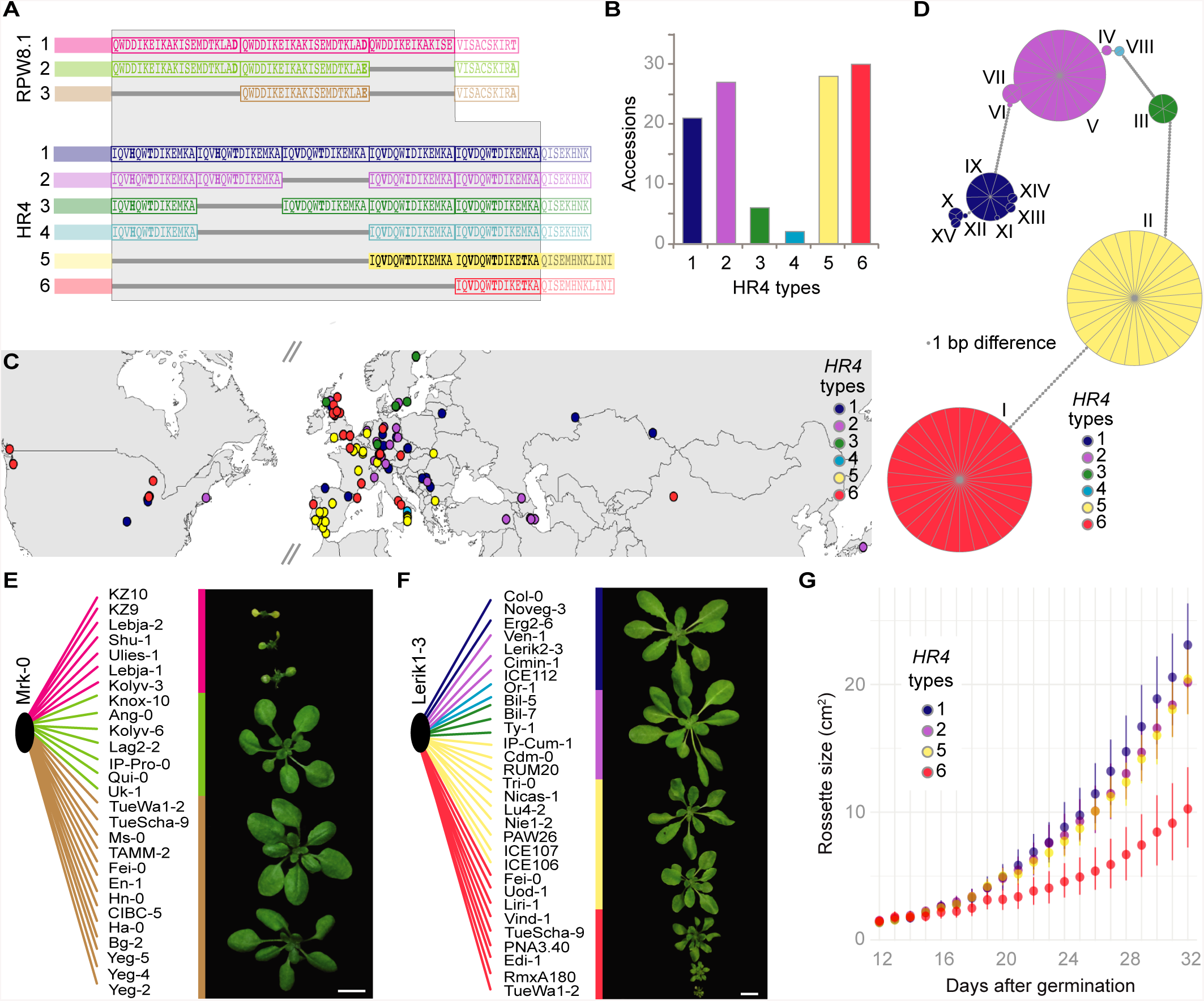
Sequence variation of a large collection of *RPW8.1* and *HR4* alleles. **(A)** Repeat polymorphisms in RPW8.1 and HR4 proteins (grey background). N-terminal regions and tails are semi-transparent. **(B)** Distribution of *HR4* types across 113 Sanger sequenced alleles (see Table S5). **(C)** Distribution of *HR4* allele types in Eurasia and North America. **(D)** Haplotype network of *HR4* alleles, with a 1-bp minimum difference. (**E)** F_1_ progeny of Mrk-0 crossed to accessions with different *RPW8.1* alleles. Short *RPW8.1* variants do not induce hybrid necrosis. **(F)** F_1_ progeny of Lerik1-3 crossed to accessions with different *HR4* alleles. The shortest *HR4* alleles (red) cause strong hybrid necrosis, the second shortest *HR4* alleles (yellow) cause mild hybrid necrosis. **(G)** Rosette growth of F_1_ progeny from Lerik1-3 and accessions carrying different *HR4* alleles. The shortest *HR4* allele causes a strong growth reduction, while the second-shortest *HR4* allele has a milder effect. Scale bars indicate 1 cm.

To confirm the short read-based length predictions, *RPW8.1* was PCR amplified from 28 accessions and *HR4* from 113 accessions (**Fig 6A-D** and **Table S5** and **S6**). This not only confirmed that the Illumina predictions were accurate, but also revealed new variants with different arrangements of *HR4* repeats, although none were as short as *HR4*^Fei-0^ or *HR4*^ICE106^ (**Fig 6A, B**). The short necrosis-risk *HR4* variants are found across much of the global range of *A. thaliana* (**Fig 6C**), whereas the much rarer necrosis-risk *RPW8.1*^KZ10^-like variant was exclusive to Central Asia. We also observed that sequences of the two short *HR4* types were more conserved than the longer ones, with each short type belonging to a single haplotype, while the long necrosis-risk *HR4* alleles belonged to multiple haplotypes (**Fig 6D**).

The extensive information on *RPW8.1/HR4* haplotypes allowed us to use test crosses to determine whether interaction with either *RPP7*^Mrk-0^ or *RPP7*^Lerik1-3^ is predictable from sequence, specifically from repeat number **(Fig 6E, F)**. As expected, accessions with the longest, Type 1, *RPW8.1*^KZ10^-like alleles **(Fig 6E, pink)** produced necrotic hybrid progeny when crossed to Mrk-0, whereas accessions carrying the two shorter Type 2 and 3 alleles did not **(Fig 6E** and **Table S7)**. The situation was similar for *HR4*; all but two of the tested accessions with the shortest *HR4*^Fei-0^-like alleles (**Fig 6F, red**) produced strongly necrotic progeny when crossed to Lerik1-3, while accessions carrying the second shortest *HR4*^ICE106^-like alleles (**Fig 6F** and **Table S8**) produced more mildly affected progeny. Hybrid progeny of Lerik1-3 and accessions carrying other *HR4* alleles did not show any signs of necrosis (**Fig 6F**). Necrosis was correlated with reduction in overall size of plants, which in turn correlated with RPW8.1/HR4 repeat length (**Fig 6F** and **Table S9**). Finally, *HR4*^Fei-0^-like alleles in two accessions caused a mild phenotype similar to *HR4*^ICE106^, suggesting the presence of genetic modifiers that partially suppress autoimmune symptoms.

## Discussion

The *RPW8*/*HR* cluster is remarkably variable in terms of copy number, reminiscent of many multi-gene clusters carrying NLR-type *R* genes [16]. While the first three genes in the cluster, *HR1, HR2* and *HR3*, are generally well conserved, there is tremendous variation in the number of the other genes in the cluster, including *RPW8.1*/*HR4*. Nevertheless, that the *HR4* hybrid necrosis-risk allele is not rare and widely distributed, accounting for half of all *HR4* carriers (**Fig 6B, C**), suggests that it might provide adaptive benefits, as postulated before for *ACD6* hybrid necrosis-risk alleles [12].

The N-terminal portion of RPW8 and HR proteins can be homology modeled on a multi-helix bundle in the animal MLKL protein [34], which in turn shares structural similarity with fungal HeLo and HELL domain proteins [37]. In both cases, the N-terminal portions can insert into membranes (with somewhat different mechanisms proposed for the two proteins), thereby disrupting membrane integrity and triggering cell death [36,46–48]. For both proteins, insertion is regulated by sequences immediately C-terminal to the multi-helix bundle [36,46–50]. It is tempting to speculate that the RPW8/HR repeats and the C-terminal tail, which together make up the C-terminal portions of the proteins, similarly regulate activity of RPW8.1 and HR4. In agreement, our chimera studies, where we exchanged and varied the number of RPW8/HR repeats and swapped the C-terminal tail, indeed point to the C-terminal portion of RPW8/HR proteins having a regulatory role. A positive regulator of RPW8-mediated disease resistance, a 14-3-3 protein, interacts specifically with the C-terminal portion of RPW8.2, consistent with this part of the protein controlling RPW8/HR activity [51]. Perhaps even more intriguing is the fact that in many fungal HeLo domains this C-terminal region is a prion-forming domain composed of 21-amino acid repeats. RPW8.1 also has 21-amino acid repeats, while HR4 has 14-amino acid repeats, although different from the fungal proteins, these are not interrupted by a spacer. In fungal HET-S and related proteins, the repeats exert regulatory function by forming amyloids and thereby causing the proteins to oligomerize [35–39]. While it remains to be investigated whether the RPW8/HR repeats and the C-terminal tail function in a similar manner, their potential regulatory function makes them a possible target for pathogen effectors. In such a scenario, at least some RPP7 proteins might act as guards for RPW8/HR proteins and sense their modification by pathogen effectors [16,52].

Can we conclude from the MLKL homology that RPW8 and HR proteins form similar pores as MLKL? Unfortunately, this is not immediately obvious, as a different mechanism has been suggested for fungal proteins with HeLo and HELL domains [35–37]. For MLKL, it has been suggested that the multi-helix bundle directly inserts into the membrane, whereas for the fungal protein, it has been proposed that the multi-helix bundle regulates the ability of an N-terminal transmembrane domain to insert into the membrane. An N-terminal transmembrane domain has been predicted for RPW8 [27], but although RPW8 proteins can be membrane associated [53,54], the insertion of this domain into the membrane has not been directly demonstrated.

We have shown that differences in protein structure, rather than expression patterns or levels, are key to the genetic interaction between RPW8/HR and RPP7. While we do not know whether the proteins interact directly, allele-specific genetic interactions are often an indicator of direct interaction between the gene products [55]. Moreover, reminiscent of RPW8/HR and RPP7 interaction, the activity of the fungal HeLo domain protein HET-S is regulated by an NLR protein [38].

In conclusion, we have described in detail an intriguing case of hybrid necrosis in *A. thaliana*, where three different pairs of alleles at a conventional complex NLR resistance gene cluster, *RPP7*, and alleles at another complex, but non-NLR resistance gene cluster, *RPW8*/*HR*, interact to trigger autoimmunity in the absence of pathogens. Our findings suggest that within the immune system, conflict does not occur randomly, but that certain pairs of loci are more likely to misbehave than others. Finally, that genes of the *RPW8*/*HR* cluster can confer broad-spectrum disease resistance, while at least one *RPP7* member can confer race-specific resistance, provides yet another link between different arms of the plant immune system [56].

## Materials and Methods

### Plant material

Stock numbers of accessions used are listed in Supplementary Material. All plants were stratified in the dark at 4°C for 4-6 days prior to planting on soil. Late flowering accessions were vernalized for six weeks under short day conditions (8 h light) at 4°C as seedlings. All plants were grown in long days (16 h light) at 16°C or 23°C at 65% relative humidity under Cool White fluorescent light of 125 to 175 μmol m^-2^ s^-1^. Transgenic seeds were selected either with 1% BASTA (Sigma-Aldrich), or by mCherry fluorescence. Constructs are listed in Table S10.

### Genotyping.by-sequencing and QTL mapping

Genomic DNA was isolated from Lerik1-3 x ICE106/ICE107 F_2_ and F_3_ individuals and from ICE79 x Don-0 F_2_ individuals using a Biosprint 96 instrument and the BioSprint 96 DNA Plant Kit (Qiagen, Hilden, Germany).The individuals represented all classes of segregating phenotypes. Genotyping-by-Sequencing (GBS) using RAD-seq was used to genotype individuals in the mapping populations with *Kpn*I tags [57]. Briefly, libraries were single-end sequenced on a HiSeq 3000 instrument (Illumina, San Diego, USA) with 150 bp reads. Reads were processed with SHORE [58] and mapped to the *A. thaliana* Col-0 reference genome. QTL was performed using R/qtl with the information from 330 individuals and 2,989 markers for the Lerik1-3 x ICE106/107 populations, and 304 individuals and 2,207 markers for the ICE79 x Don-0 population. The severity of the hybrid phenotype was scored as a quantitative trait.

### GWAS

Lerik1-3-dependent hybrid necrosis in F_1_ progeny from crosses with 80 accessions [10] was scored as 1 or 0. The binary trait with accession information was submitted to the easyGWAS platform [40], using the FaSTLMM algorithm. A -log_10_(p-value) was calculated for every SNP along the five *A. thaliana* chromosomes.

### *RPW8.1*/*HR4* length prediction

Short reads from the 1001 Genomes project (http://1001genomes.org) were mapped using SHORE[58] with 5 mismatches allowed per read. Sequences of the *RPW8*/*HR* clusters from Col-0 and KZ10 were provided as references and the covered region for *RPW8.1*^KZ10^ and *HR4*^Col-0^ was retrieved.

### *RPW8.1*/*HR4* sequence analysis

Overlapping fragments covering the *HR4/RPW8.1* genomic region were PCR amplified from different *A. thaliana* accessions (oligonucleotides in Table S11). Fragments were cloned and Sanger sequenced. A maximum-likelihood tree of coding portions of exons and introns was computed using RaxML [59] and visualized with Figtree.

### Population genetic analysis

The geographical distribution of the 113 accessions carrying different *HR4* alleles was plotted using R (version 0.99.903). Packages maps, mapdata, mapplots and scales were used. A haplotype network was built using a cDNA alignment of 113 *HR4* alleles from different accessions. The R packages used were ape (dist.dna function) and pegas (haploNet function).

### Histology

Cotyledons from 18 day-old seedlings were collected and 1 ml of lactophenol Trypan Blue solution (20 mg Trypan Blue, 10 g phenol, 10 ml lactic acid, 10 ml glycerol and 10 ml water) diluted 1 : 2 in 96% ethanol was added for 1 hour at 70°C. Trypan Blue was removed, followed by the addition of 1 ml 2.5g/ml chloral hydrate and an overnight incubation. The following day, the de-stained cotyledons were transferred to 50% glycerol and mounted on slides.

### Oligonucleotides

See Table S11.

### Data availability

DNA sequences have been deposited with GenBank under accession numbers MK598747 and MK604929-MK604934.

## Acknowledgements

We thank Jane Parker for the *H. arabidopsidis* Hiks1 isolate, Katrin Fritschi and Camilla Kleinhempel for technical support, Gautam Shirsekar for help with the pathology assays and discussions, and Christian Kubica for help with visualization of PacBio sequence data.

## Author contributions

**Conceptualization:** CB, DW, EC.

**Data curation:** EC.

**Formal analysis:** CB, STKJH, ALVdW, IB, GW, EC.

**Funding acquisition:** RW, DW, EC. **Investigation:** CB, RW, AH, WX, MZ, WWHH, EC. **Methodology:** CB, EC.

**Project administration:** DW. **Supervision:** DW. **Validation:** CB, EC.

**Writing – original draft:** CB, EC.

**Writing – review & editing:** CD, DW, EC.

## Supplemental Information

### Supplemental Figures

**Fig S1.**
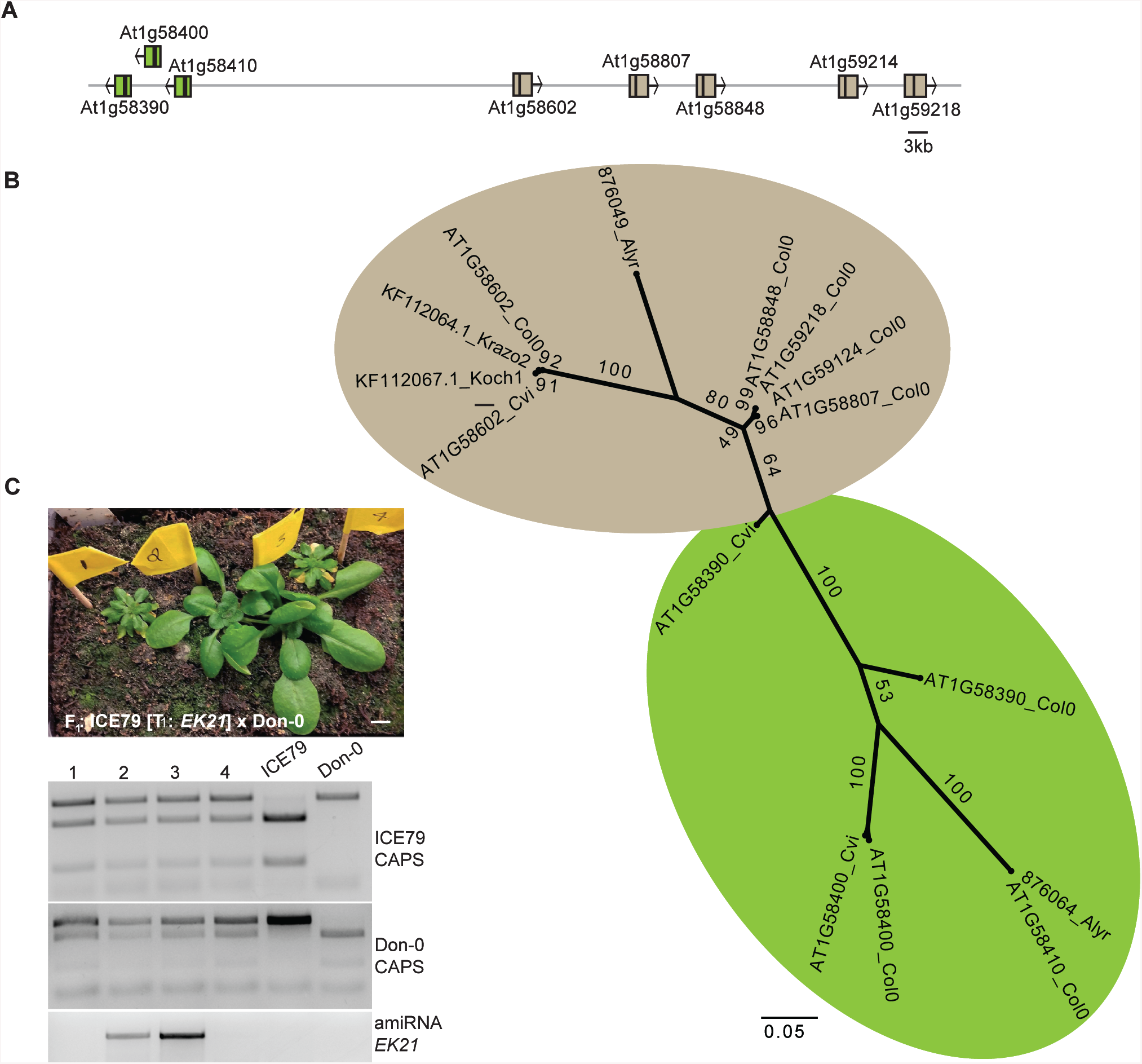
Role of the *RPP7* cluster in *DM6-DM7* dependent hybrid necrosis. Related to Fig 1. **(A)** *RPP7* cluster in the Col-0 reference genome. The left portion of the cluster consists of three *NLR* genes, *At1g58390, At1g58400* and *At1g58410* (green arrows). The right portion includes five *NLR* genes, *At1g58602, At1g58807, At1g58848, At1g59214* and *At1g59218* (brown arrows). Twenty-two non-NLR genes in this region are not shown. **(C)** Maximum-likelihood tree of *NLR* genes in the *RPP7* cluster based on the NB domain. *At1g59124* and *At1g58807* sequences are identical, as are *At1g59218* and *At1g58848.* Same colors as in (A). Bootstrap values (out of 100) are indicated on each branch. **(C)** Representative rescue experiment using an amiRNA construct targeting *RPP7* homologs (see Table S1). ICE79 was transformed with the amiRNA construct *EK21* and T_1_ plants were crossed to Don-0, resulting in rescued and non-rescued plants segregating in the F_1_ progeny. Parental genotypes were confirmed with CAPS markers, shown below. Five-week old plants grown in 16°C are shown.

**Fig S2.**
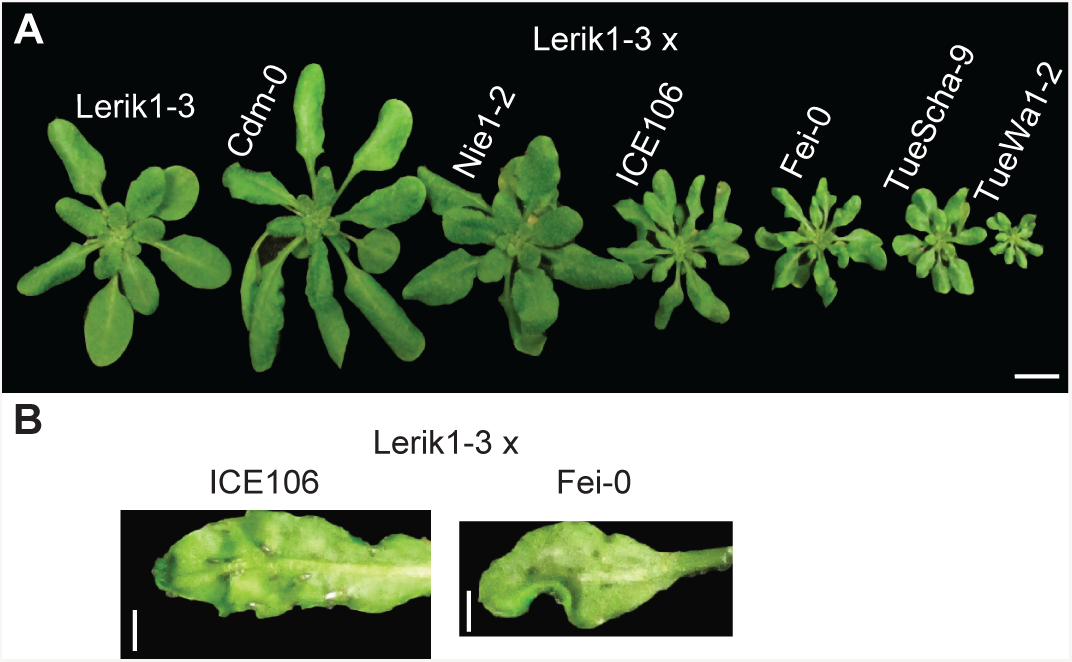
Phenotypic variation in Lerik1-3 F_1_ hybrids. Related to Fig 1. Major differences were observed in rosette size of F_1_ hybrids **(A)** and spotted cell death on the abaxial side of leaves **(B)**. Scale bar represents 1cm **(A)** and 1mm **(B)**. Plants were five weeks old.

**Fig S3.**
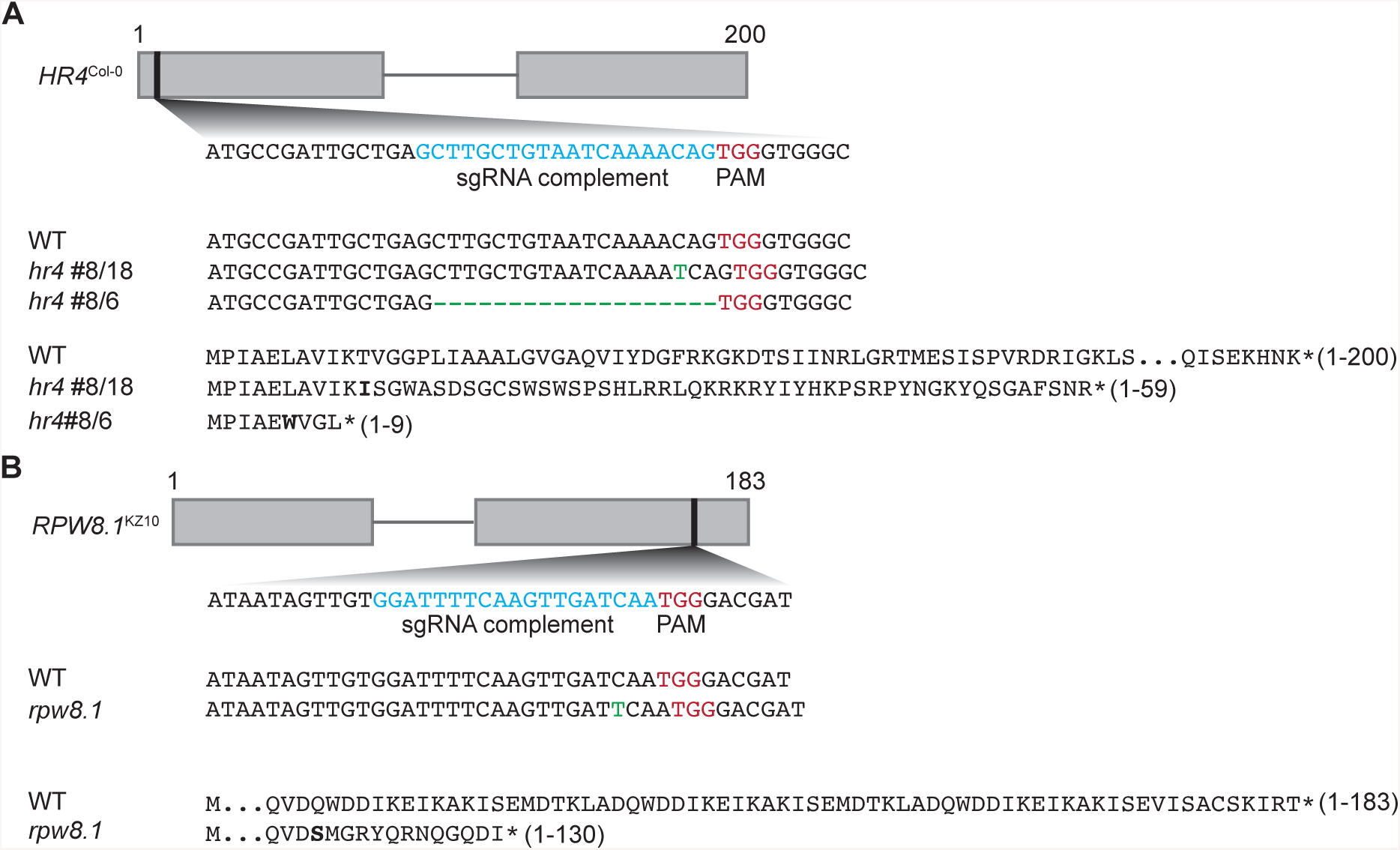
*HR4* and *RPW8.1* CRISPR/Cas9 knockout lines. Related to Fig 3 and Fig S4. **(A)** Two alleles of *HR4* in Col-0 with a 1-bp insertion (#8/18) or a 19-bp deletion (#8/6) were identified by amplicon sequencing. **(B)** An allele of *RPW8.1* in KZ10 with a 1-bp insertion was recovered. The stop codons are marked with an asterisk and the first amino acid after a frameshifting event is in bold.

**Fig S4.**
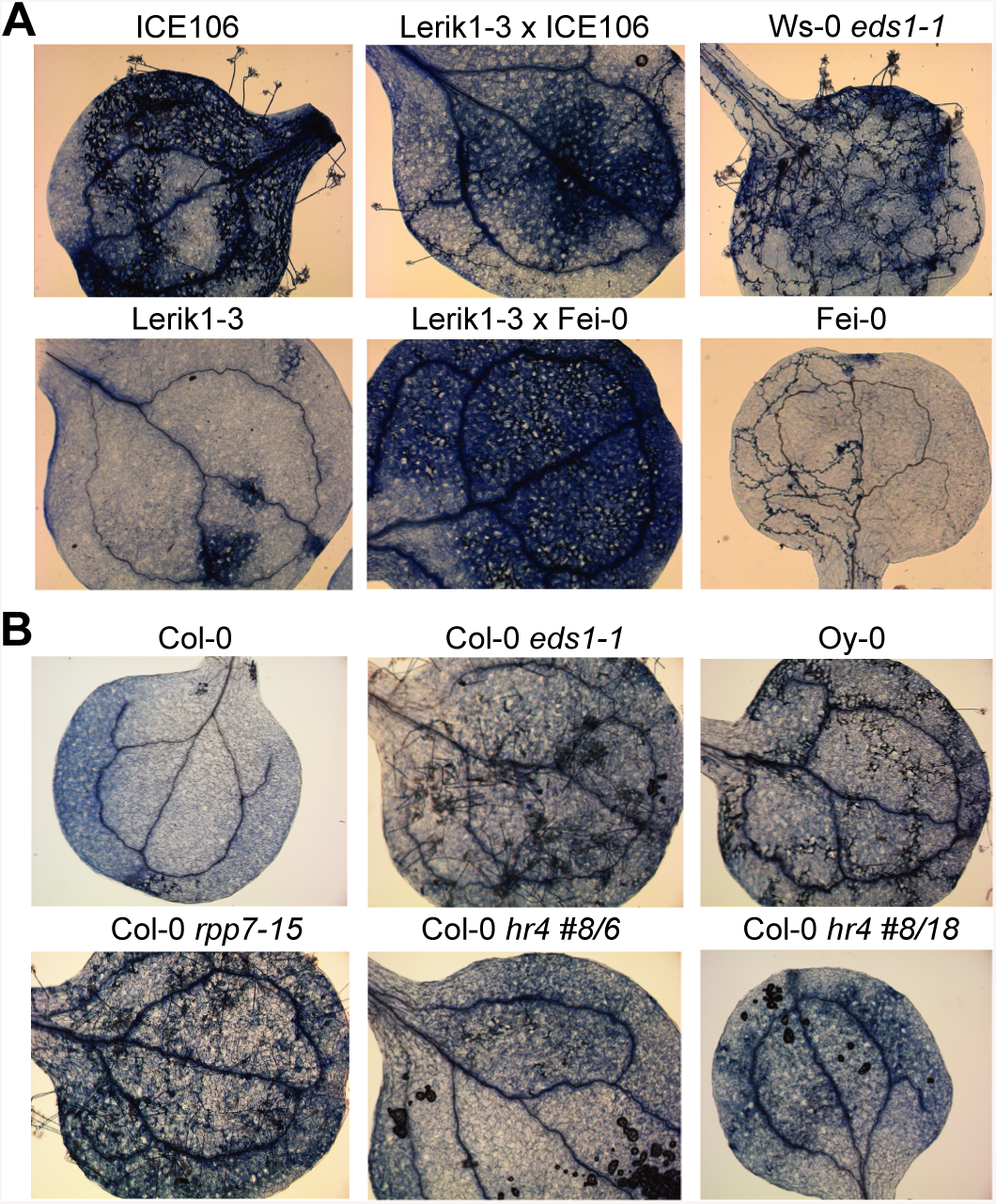
Resistance and susceptibility to *H. arabidopsidis* isolate Hiks1. **(A)** Trypan Blue stained cotyledons 5 days after infection. Lerik1-3 is resistant, while Fei-0 and ICE106 are fully susceptible. The F_1_ hybrids Lerik1-3 x Fei-0 and Lerik1-3 x ICE106 appear to be less resistant than Lerik1-3. Ws-0 *eds1-1* is a positive infection control. **(B)** Two different *hr4* loss-of-function alleles (see Fig S3) are as resistant as Col-0 wild-type plants. *eds1-1* and *rpp7-15* are positive infection controls.

**Fig S5.**
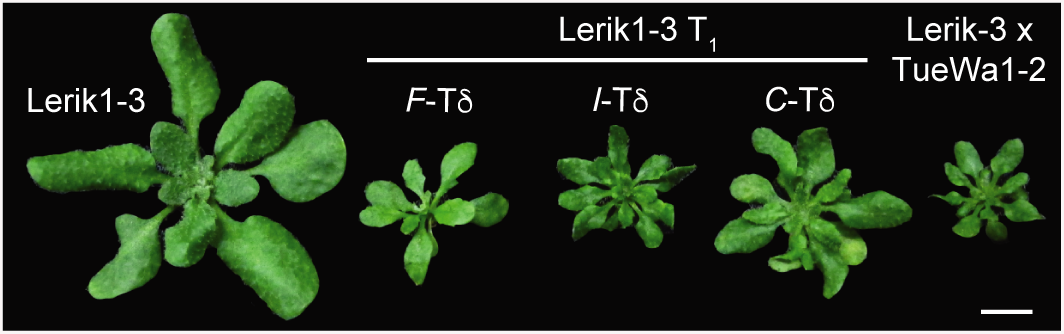
Hybrid necrosis by introduction of chimeras. Related to Fig 5. Effects of chimeric *HR4* transgenes introduced into Lerik1-3, with negative and positive controls shown to the left and right. Scale bar represents 1cm. Five week-old plants are shown.

**Fig S6.**
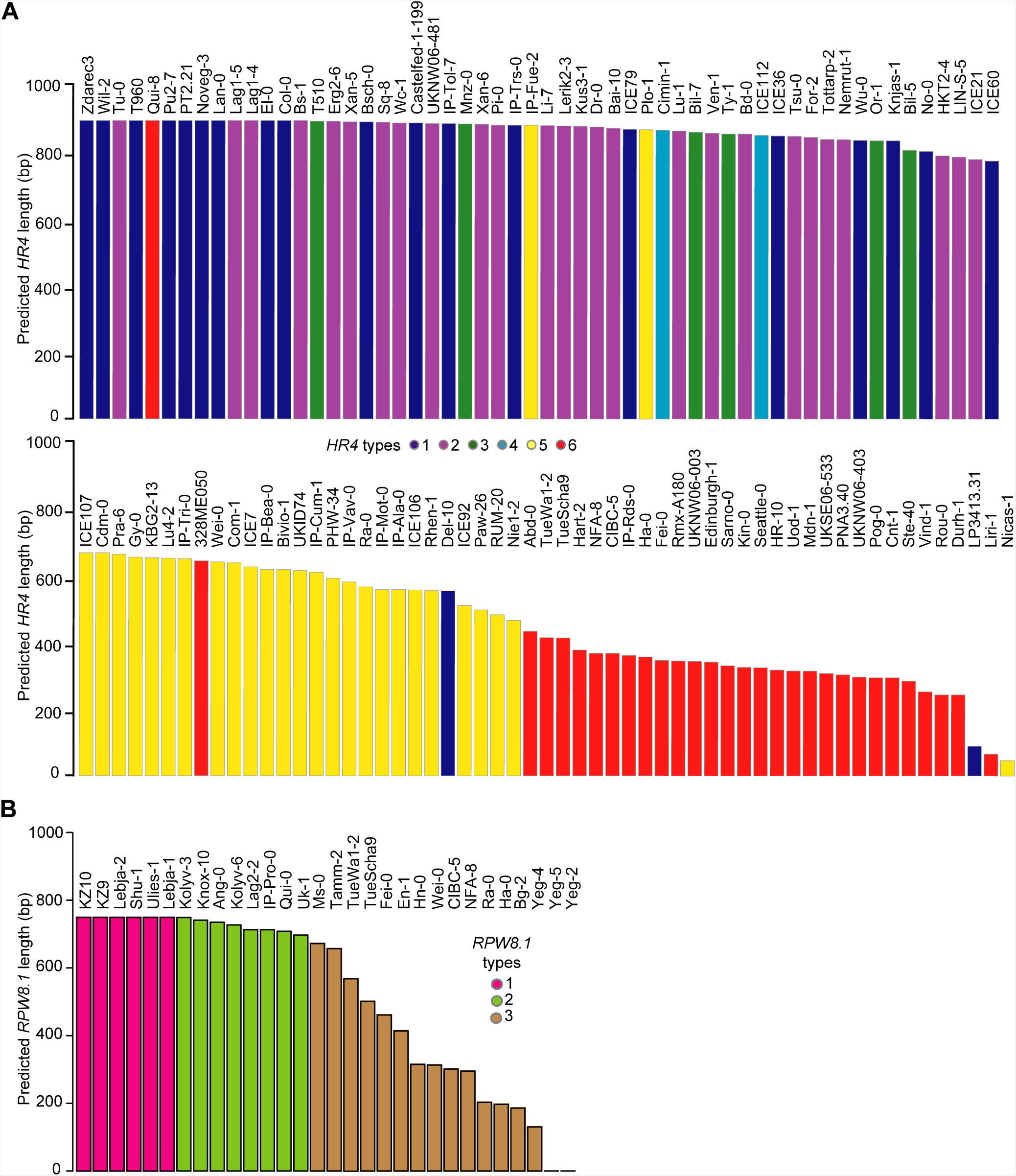
Predicted lengths of *HR4* and *RPW8.1* coding sequences from remapping of short reads from the 1001 Genomes Project. Related to Fig 6. **(A)** *HR4* type assignments based on information from Sanger sequencing. **(B)** *RPW8.1* type based on information from Sanger sequencing.

### Supplemental Tables

**Table S1.**
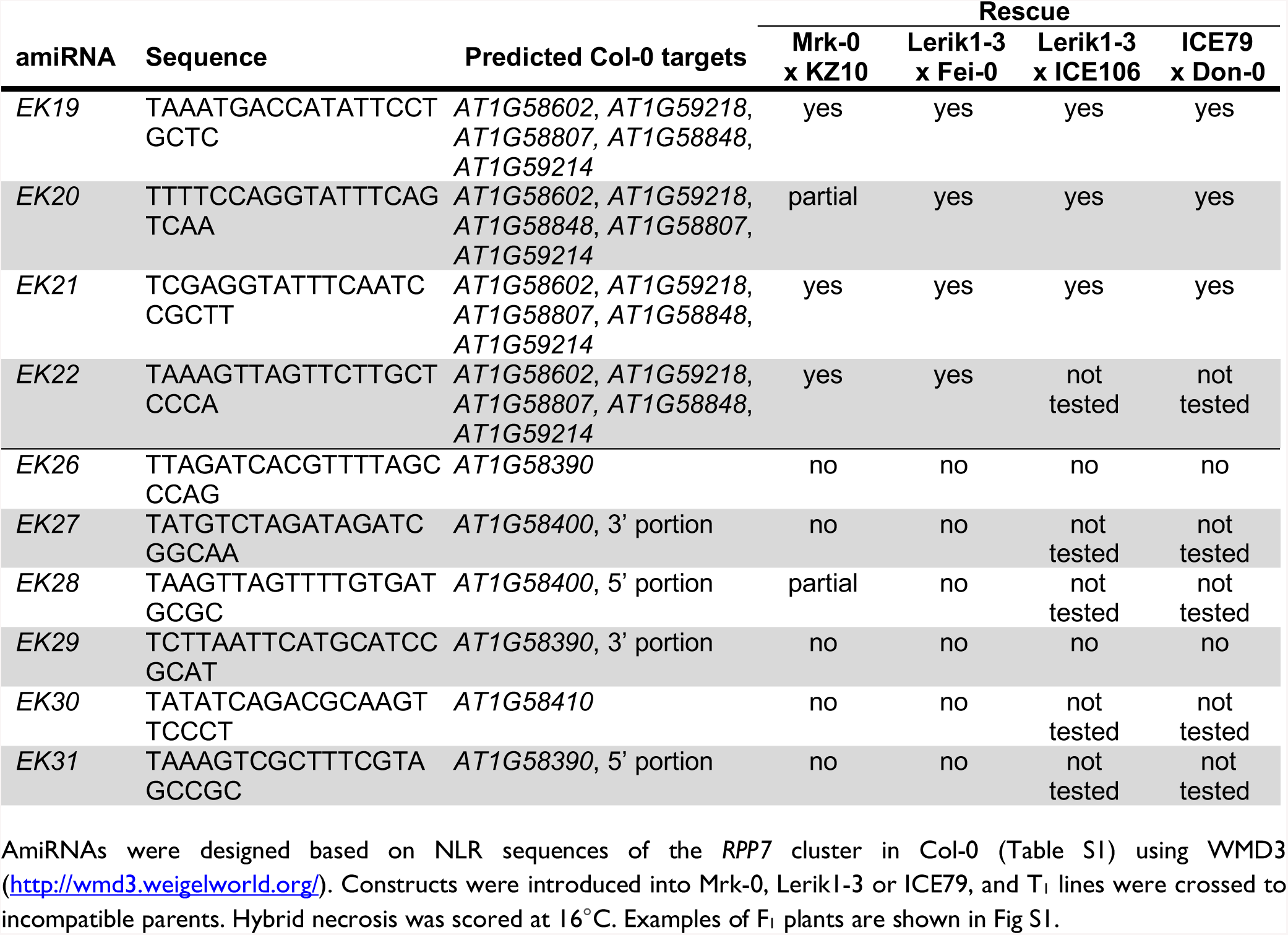
Rescue of hybrid necrosis by amiRNAs against *RPP7* homologs. Related to Fig 1.

**Table S2.**
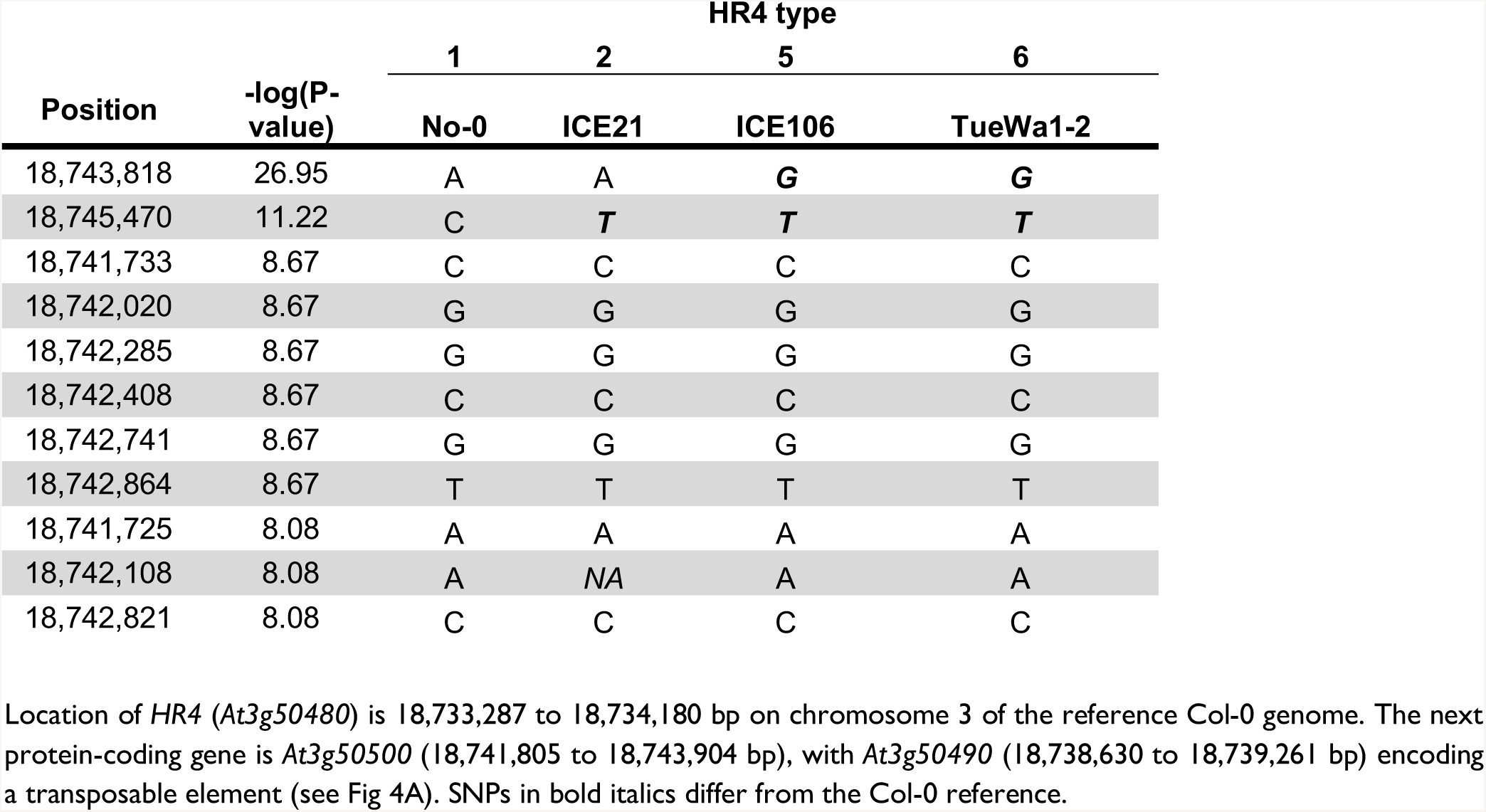
GWAS hits on chromosome 3 from Lerik1-3 x 80 accessions panel and tagging SNPs present in accessions carrying different HR4 types. Related to Fig 2.

**Table S3.**
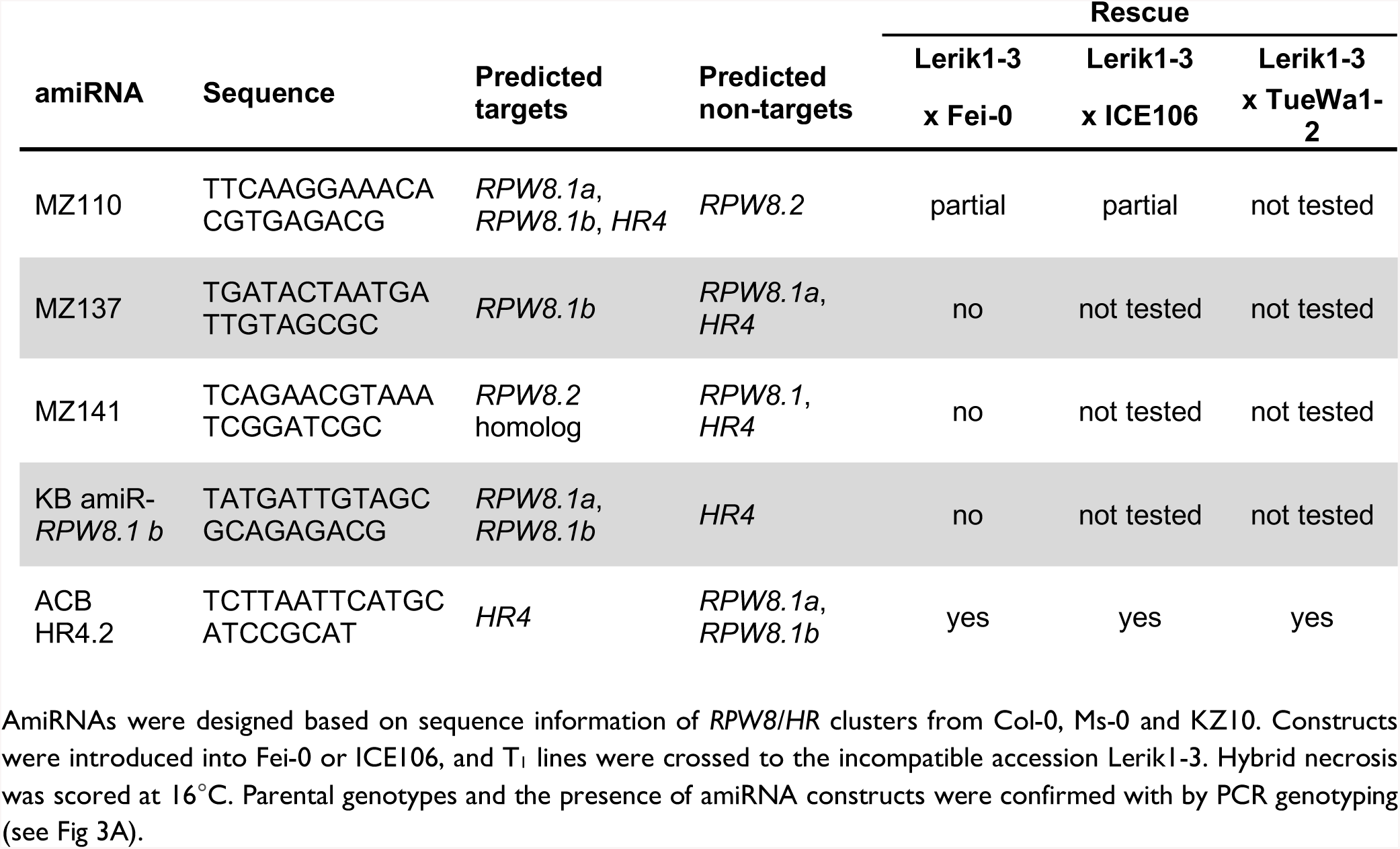
Rescue effects of amiRNAs targeting *RPW8* homologs. Related to Fig 1 and Fig 3.

**Table S4.**
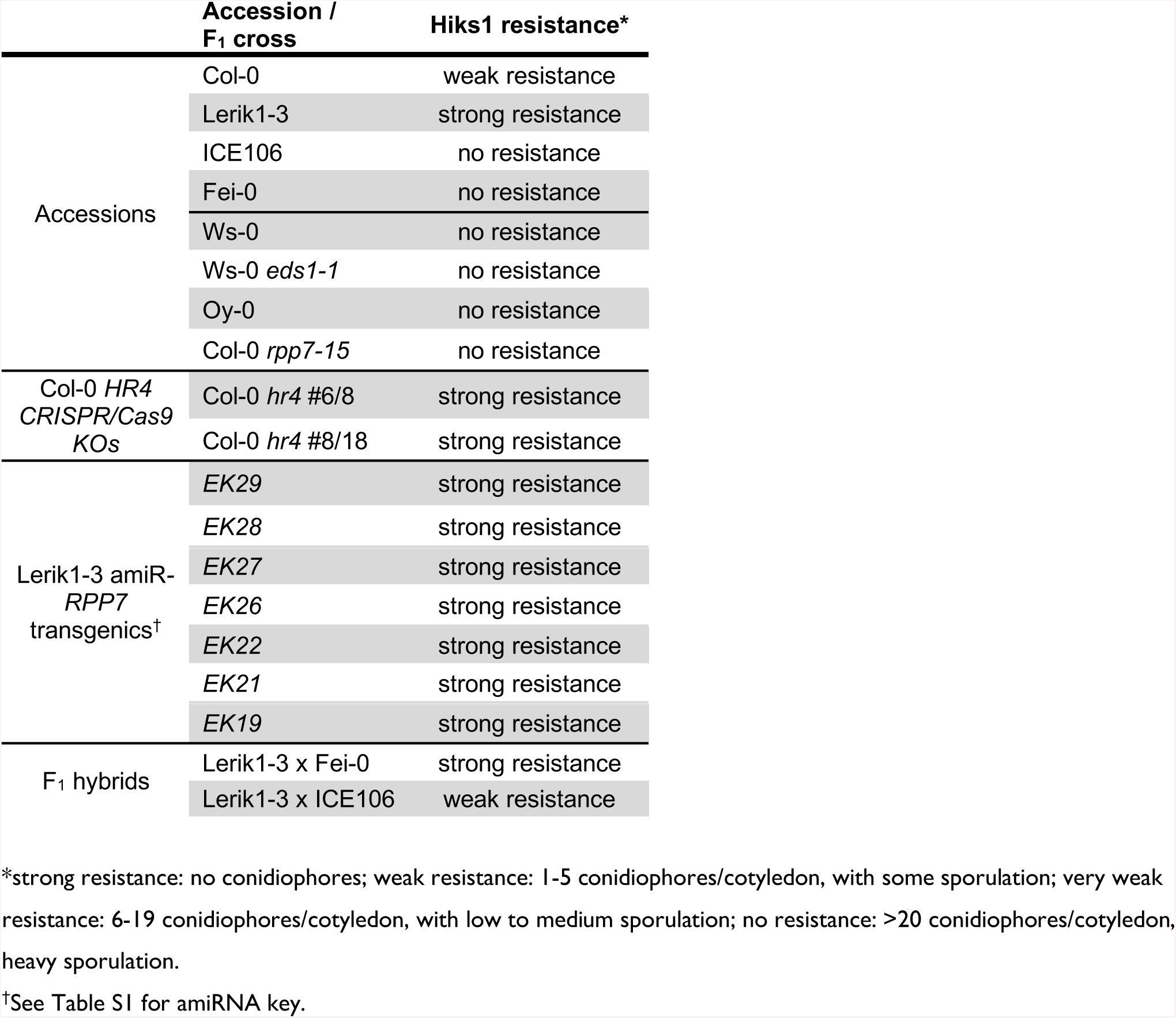
Resistance to the *H. arabidopsidis* isolate Hiks1. Related to Fig S4.

**Table S5.** Accessions for *HR4* survey. Related to Fig 6.

| Accession | 1001 Genomes<br>Project ID | Stock<br>center ID | *HR4*<br>type | *HR4*<br>haplotype | Covered<br>region (bp) |
| --- | --- | --- | --- | --- | --- |
| IP-Tol-7 | 9588 | CS77371 | 1 | XV | 885 |
| IP-Trs-0 | 9590 | CS77387 | 1 | XV | 880 |
| El-0 | 7117 | CS76479 | 1 | XIV | 894 |
| Wu-0 | 7415 | CS78858 | 1 | XIV | 835 |
| Castelfed-1-199 | 9683 | CS76748 | 1 | XIII | 887 |
| ICE79/Voeran-1 | 9979 | CS76352 | 1 | XIII | 868 |
| T960 | 6148 | CS77325 | 1 | XII | 894 |
| ICE60/Stepn-2 | 9955 | CS76377 | 1 | XI | 773 |
| ICE36/Dobra-1 | 10018 | CS76369 | 1 | X | 848 |
| No-0 | 7273 | CS77128 | 1 | X | 802 |
| Knjas-1 | 9749 | CS76971 | 1 | X | 834 |
| LP3413.31 | 8464 | CS79030 | 1 | IX | 894 |
| Zdarec3 | 403 | CS78873 | 1 | IX | 894 |
| Del-10 | 10016 | CS76397 | 1 | IX | 554 |
| Lan-0 | 7208 | CS76539 | 1 | IX | 894 |
| Col-0 | 6909 | CS76539 | 1 | IX | 894 |
| Noveg-3 | 9638 | CS77133 | 1 | IX | 894 |
| Pu2-7 | 6956 | CS76580 | 1 | IX | 894 |
| Wil-2 | 7413 | CS78856 | 1 | IX | 894 |
| PT2.21 | 8077 | CS77191 | 1 | IX | 894 |
| Bsch-0 | 7031 | CS76457 | 1 | IX | 890 |
| Wc-1 | 7404 | CS76627 | 2 | V | 887 |
| Ven-1 | 7384 | CS76624 | 2 | V | 856 |
| UKNW06-481 | 5644 | CS78798 | 2 | V | 885 |
| Tu-0 | 7375 | CS76617 | 2 | V | 894 |
| ICE21/Petro-1 | 10017 | CS76370 | 2 | V | 778 |
| Lu-1 | 8334 | CS77056 | 2 | V | 863 |
| LIN-S-5 | 915 | CS77040 | 2 | V | 785 |
| Tsu-0 | 7373 | CS77389 | 2 | V | 847 |
| Bd-0 | 7013 | CS76445 | 2 | V | 854 |
| Bai-10 | 9779 | CS76682 | 2 | V | 871 |
| Kus3-1 | 9802 | CS76991 | 2 | V | 877 |
| Lag1-4 | 9102 | CS76999 | 2 | IV | 894 |
| Lag1-5 | 9103 | CS77000 | 2 | IV | 894 |
| HKT2-4 | 9995 | CS76404 | 2 | V | 789 |
| Dr-0 | 7106 | CS78897 | 2 | V | 875 |
| Pi-0 | 7298 | CS76572 | 2 | V | 880 |
| For-2 | 5741 | CS78783 | 2 | V | 844 |
| Erg2-6 | 9784 | CS76845 | 2 | V | 892 |
| Bay-0 | 6899 | CS22633 | 2 | V | - |
| Sq-8 | 6967 | CS76604 | 2 | V | 889 |
| Li-7 | 7231 | CS77035 | 2 | VI | 879 |
| Bs-1 | 7003 | CS78888 | 2 | V | 894 |
| Tottarp-2 | 6243 | CS77381 | 2 | V | 838 |
| Xan-6 | 9070 | CS78862 | 2 | VII | 883 |
| Nemrut-1 | 9993 | CS76398 | 2 | VII | 837 |
| Lerik2-3 | 9081 | CS77026 | 2 | VII | 878 |
| Xan-5 | 9069 | CS78861 | 2 | VII | 890 |
| Ty-1 | 7351 | CS78790 | 3 | III | 854 |
| Or-1 | 6074 | CS77150 | 3 | III | 834 |
| Bil-5 | 6900 | CS76709 | 3 | III | 805 |
| T510 | 6109 | CS77301 | 3 | III | 892 |
| Mnz-0 | 7244 | CS76552 | 3 | III | 884 |
| Bil-7 | 6901 | CS76710 | 3 | III | 859 |
| Cimin-1 | 9661 | CS76771 | 4 | VIII | 865 |
| ICE112/Moran-1 | 9967 | CS76427 | 4 | VIII | 850 |
| Bivio-1 | 9649 | CS76713 | 5 | II | 618 |
| Cdm-0 | 9943 | CS76410 | 5 | II | 668 |
| IP-Mot-0 | 9560 | CS77109 | 5 | II | 558 |
| Nicas-1 | 9658 | CS77127 | 5 | II | 46 |
| Wei-0 | 6979 | CS76628 | 5 | II | 641 |
| Rhen-1 | 7316 | CS78916 | 5 | II | 555 |
| ICE106/Mammo-1 | 9964 | CS76365 | 5 | II | 557 |
| ICE92/Angit-1 | 9981 | CS76366 | 5 | II | 510 |
| IP-Bea-0 | 9522 | CS76695 | 5 | II | 618 |
| IP-Ala-0 | 9515 | CS76650 | 5 | II | 558 |
| IP-Cum-1 | 9537 | CS76787 | 5 | II | 610 |
| Paw-26 | 2171 | CS77164 | 5 | II | 497 |
| IP-Vav-0 | 9511 | CS78835 | 5 | II | 581 |
| KBG2-13 | 9770 | CS76966 | 5 | II | 653 |
| ICE107/Mammo-2 | 9965 | CS76364 | 5 | II | 668 |
| Com-1 | 7092 | CS76469 | 5 | II | 638 |
| UKID74 | 5779 | CS78789 | 5 | II | 615 |
| PHW-34 | 8244 | CS77174 | 5 | II | 592 |
| PLO-1 | 9923 | CS77180 | 5 | II | 867 |
| IP-Tri-0 | 9900 | CS77386 | 5 | II | 650 |
| ICE7/Lecho-1 | 9987 | CS76371 | 5 | II | 626 |
| RUM-20 | 9925 | CS77226 | 5 | II | 483 |
| Lu4-2 | 9792 | CS77058 | 5 | II | 652 |
| Gy-0 | 8214 | CS78901 | 5 | II | 655 |
| Ra-0 | 6958 | CS76582 | 5 | II | 566 |
| IP-Fue-2 | 9541 | CS76871 | 5 | II | 880 |
| Nie1-2 | 9996 | CS76402 | 5 | II | 466 |
| Pra-6 | 9948 | CS76416 | 5 | II | 664 |
| 328ME059 | 8584 | CS76641 | 6 | I | 644 |
| Abd-0 | 6986 | CS76429 | 6 | I | 433 |
| CIBC-5 | 6908 | CS78894 | 6 | I | 367 |
| Cnt-1 | 5726 | CS78782 | 6 | I | 294 |
| Durh-1 | 7107 | CS76477 | 6 | I | 243 |
| Edinburgh-1 | 9298 | CS76832 | 6 | I | 341 |
| Fei-0 | 9941 | CS76412 | 6 | I | 346 |
| Ha-0 | 7163 | CS76500 | 6 | I | 356 |
| Hart-2 | 9799 | CS76913 | 6 | I | 377 |
| HR-10 | 6923 | CS76940 | 6 | I | 317 |
| IP-Rds-0 | 9573 | CS77206 | 6 | I | 361 |
| Kin-0 | 6926 | CS76527 | 6 | I | 325 |
| Liri-1 | 9654 | CS77041 | 6 | I | 65 |
| Mdn-1 | 1829 | CS77077 | 6 | I | 314 |
| NFA-8 | 6944 | CS78913 | 6 | I | 367 |
| PNA3.40 | 7947 | CS77184 | 6 | I | 303 |
| Pog-0 | 7306 | CS76576 | 6 | I | 294 |
| QUI-8 | 9934 | CS77199 | 6 | I | 894 |
| Rmx-A180 | 7525 | CS77218 | 6 | I | 344 |
| Rou-0 | 7320 | CS76591 | 6 | I | 243 |
| Sarno-0 | 9660 | CS77236 | 6 | I | 330 |
| Seattle-0 | 7332 | CS76598 | 6 | I | 324 |
| Ste-40 | 2317 | CS77278 | 6 | I | 284 |
| TueScha-9 | 10000 | CS76401 | 6 | I | 413 |
| TueWa1-2 | 10002 | CS76405 | 6 | I | 414 |
| UKNW06-003 | 5353 | CS78792 | 6 | I | 343 |
| UKNW06-403 | 5577 | CS78797 | 6 | I | 296 |
| UKSE06-533 | 5276 | CS78806 | 6 | I | 307 |
| Uod-1 | 6975 | CS76621 | 6 | I | 314 |
| Vind-1 | 7387 | CS76625 | 6 | I | 252 |

Covered region indicates the length of *HR4*<sup>Col-0</sup> (894 bp) covered by reads from the 1001 Genomes Project (http://1001genomes.org), allowing for five mismatches. *HR4* types are categorized according to the number of *RPW8/HR* repeats, and the haplotype is based on the entire *HR4* coding sequence.

**Table S6.** Accessions for *RPW8.1* survey. Related to Fig 6.

| Accession | 1001 Genomes<br>Project ID | Stock<br>center ID | *RPW8.1*<br>type | Covered<br>region (bp) |
| --- | --- | --- | --- | --- |
| KZ-10 | 10019 | CS28435 | 1 | 749 |
| KZ-9 | 6931 | CS76537 | 1 | 749 |
| Lebja-2 | 9632 | CS77016 | 1 | 749 |
| Shu-1 | 14318 | CS78930 | 1 | 749 |
| Ulies-1 | 9737 | CS78815 | 1 | 749 |
| Lebja-1 | 9631 | CS77015 | 1 | 749 |
| Kolyv-3 | 9626 | CS76978 | 1 | 749 |
| Knox-10 | 6927 | CS76973 | 2 | 741 |
| Ang-0 | 6992 | CS76436 | 2 | 735 |
| Kolyv-6 | 9628 | CS76980 | 2 | 727 |
| Lag2-2 | 9990 | CS76390 | 2 | 713 |
| IP-Pro-0 | 9571 | CS78914 | 2 | 713 |
| Qui-0 | 9949 | CS76417 | 2 | 708 |
| Uk-1 | 7378 | CS76620 | 2 | 697 |
| Ms-0 | 6938 | CS76555 | 3 | 672 |
| Tamm-2 | 6968 | CS76610 | 3 | 657 |
| TueWa1-2 | 10002 | CS76405 | 3 | 568 |
| TueScha-9 | 10000 | CS76401 | 3 | 501 |
| Fei-0 | 9941 | CS76412 | 3 | 461 |
| En-1 | 8290 | CS76841 | 3 | 414 |
| Hn-0 | 7165 | CS76513 | 3 | 315 |
| CIBC-5 | 6908 | CS78894 | 3 | 301 |
| NFA-8 | 6944 | CS78913 | 3 | 295 |
| Ha-0 | 7163 | CS76500 | 3 | 197 |
| Bg-2 | 6709 | CS28069 | 3 | 186 |
| Yeg-4 | 9130 | CS78865 | 3 | 130 |
| Yeg-5 | 9131 | CS78866 | 3 | 0 |
| Yeg-2 | 9128 | CS78864 | 3 | 0 |

**Table S7.**
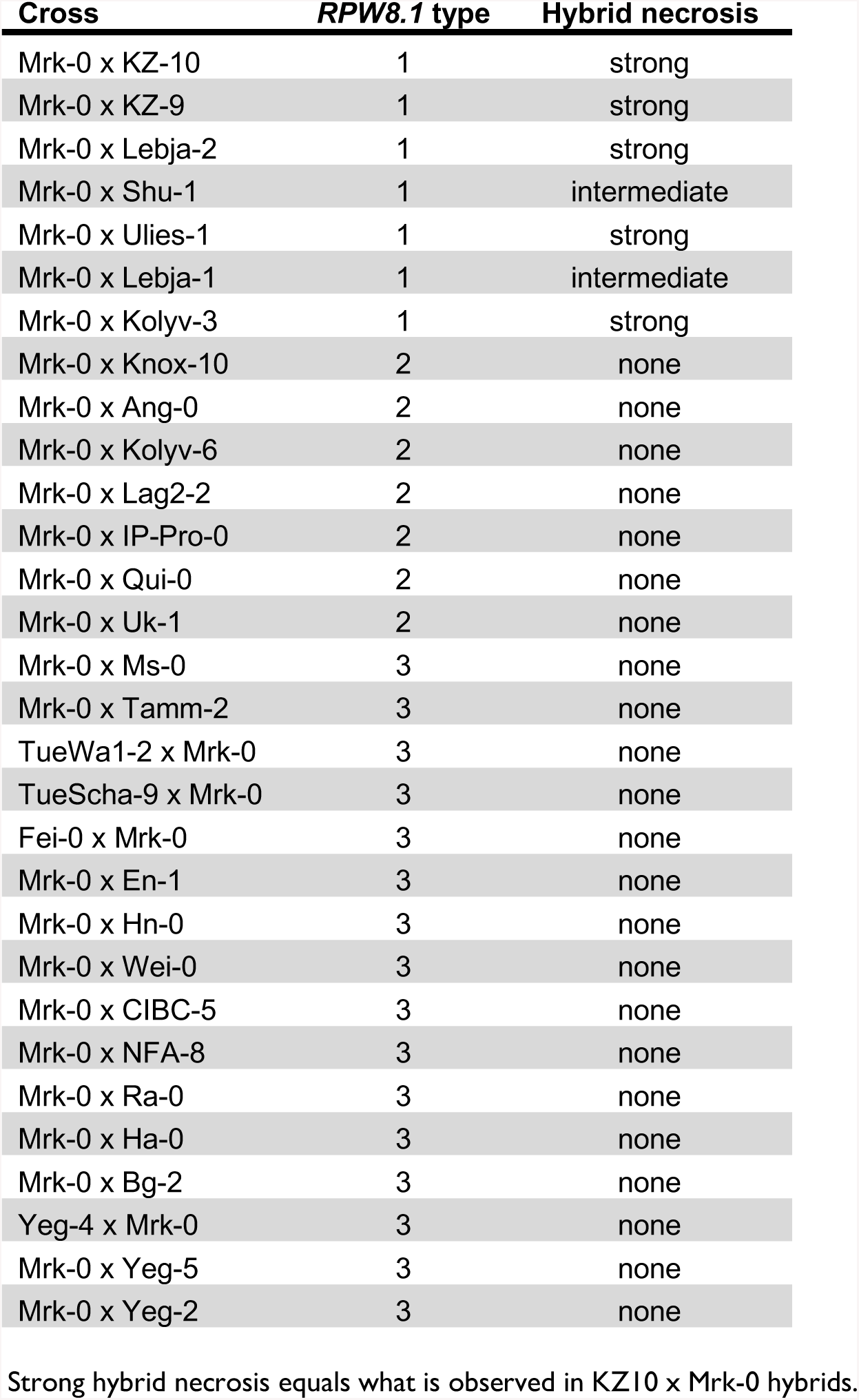
Hybrid necrosis in F_1_ plants of Mrk-0 crossed to other accessions. Related to Fig 6.

**Table S8.** Hybrid necrosis in F_1_ plants of Lerik1-3 crossed to other accessions. Related to Fig 6.

| **Cross** | **HR4 type** | **Hybrid necrosis** |
| --- | --- | --- |
| Lerik1-3 x Col-0 | 1 | none |
| Lerik1-3 x Noveg-3 | 1 | none |
| Lerik1-3 x Lerik2-3 | 2 | none |
| Lerik1-3 x Ven-1 | 2 | none |
| Ty-1 x Lerik1-3 | 3 | none |
| Lerik1-3 x Bil-5 | 3 | none |
| Lerik1-3 x Bil-7 | 3 | none |
| Lerik1-3 x Or-1 | 3 | none |
| Lerik1-3 x ICE112 | 4 | none |
| Lerik1-3 x Cimin-1 | 4 | none |
| Lu4-2 x Lerik1-3 | 5 | mild |
| Lerik1-3 x Lu4-2 | 5 | mild |
| Lerik1-3 x Tri-0 | 5 | mild |
| Nicas-1 x Lerik1-3 | 5 | mild |
| Lerik1-3 x IP-Cum | 5 | mild |
| Lerik1-3 x RUM20 | 5 | mild |
| Lerik1-3 x PAW26 | 5 | mild |
| Uod-1 x Lerik1-3 | 6 | intermediate |
| Lerik1-3 x Liri-1 | 6 | intermediate |
| Lerik1-3 x Vind-1 | 6 | strong |
| Ste-40 x Lerik1-3 | 6 | strong |
| Pog-0 x Lerik1-3 | 6 | strong |
| Lerik1-3 x RmxA180 | 6 | strong |
| Lerik1-3 x Edinburgh-1 | 6 | strong |
| Lerik1-3 x PNA3.40 | 6 | strong |

Strong hybrid necrosis equals what is observed in Lerik1-3 x Fei-0 F<sub>1</sub> hybrids.

**Table S9.**
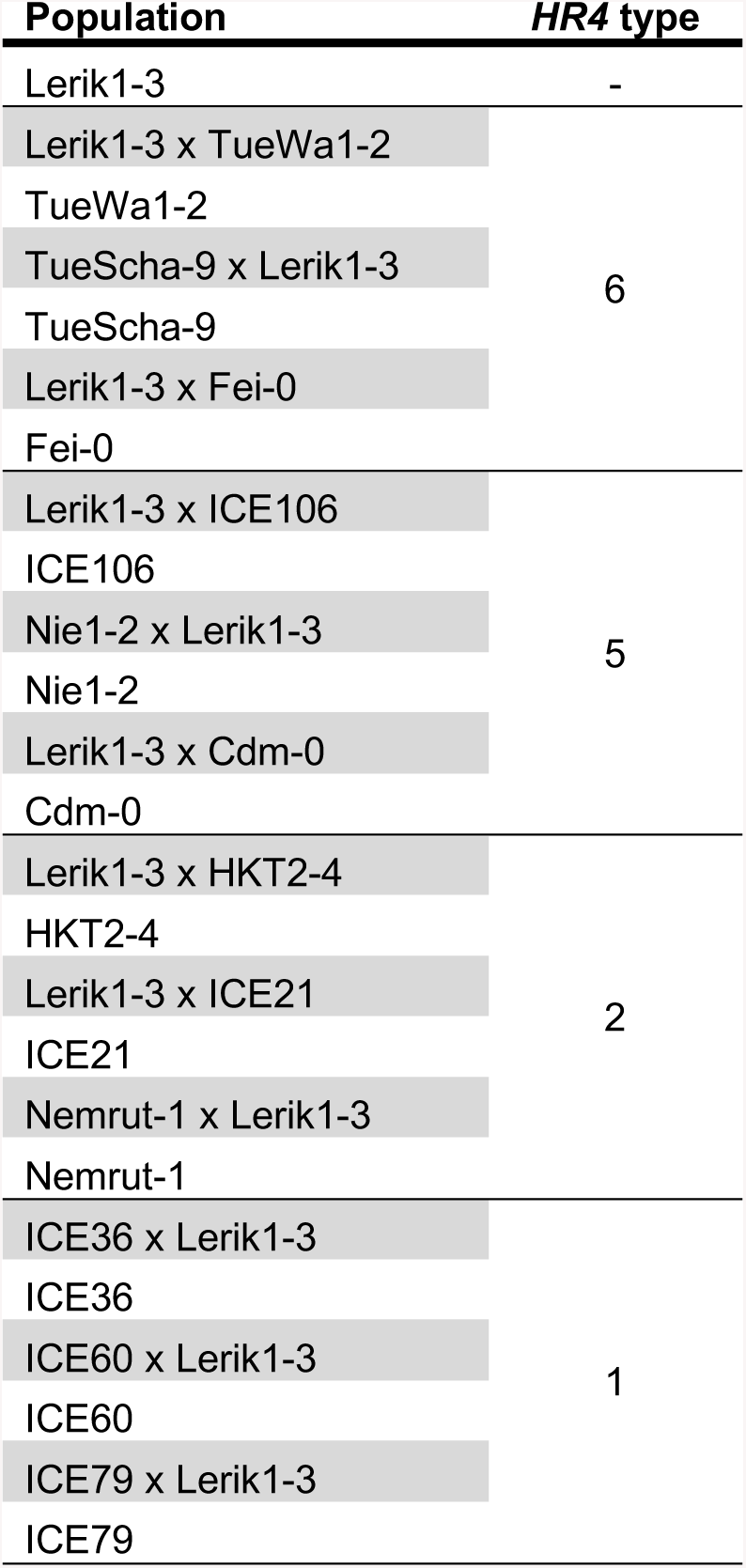
Accessions and hybrids in which growth was analyzed with the automated phenotyping platform RAPA. Related to Fig 6.

**Table S10.**
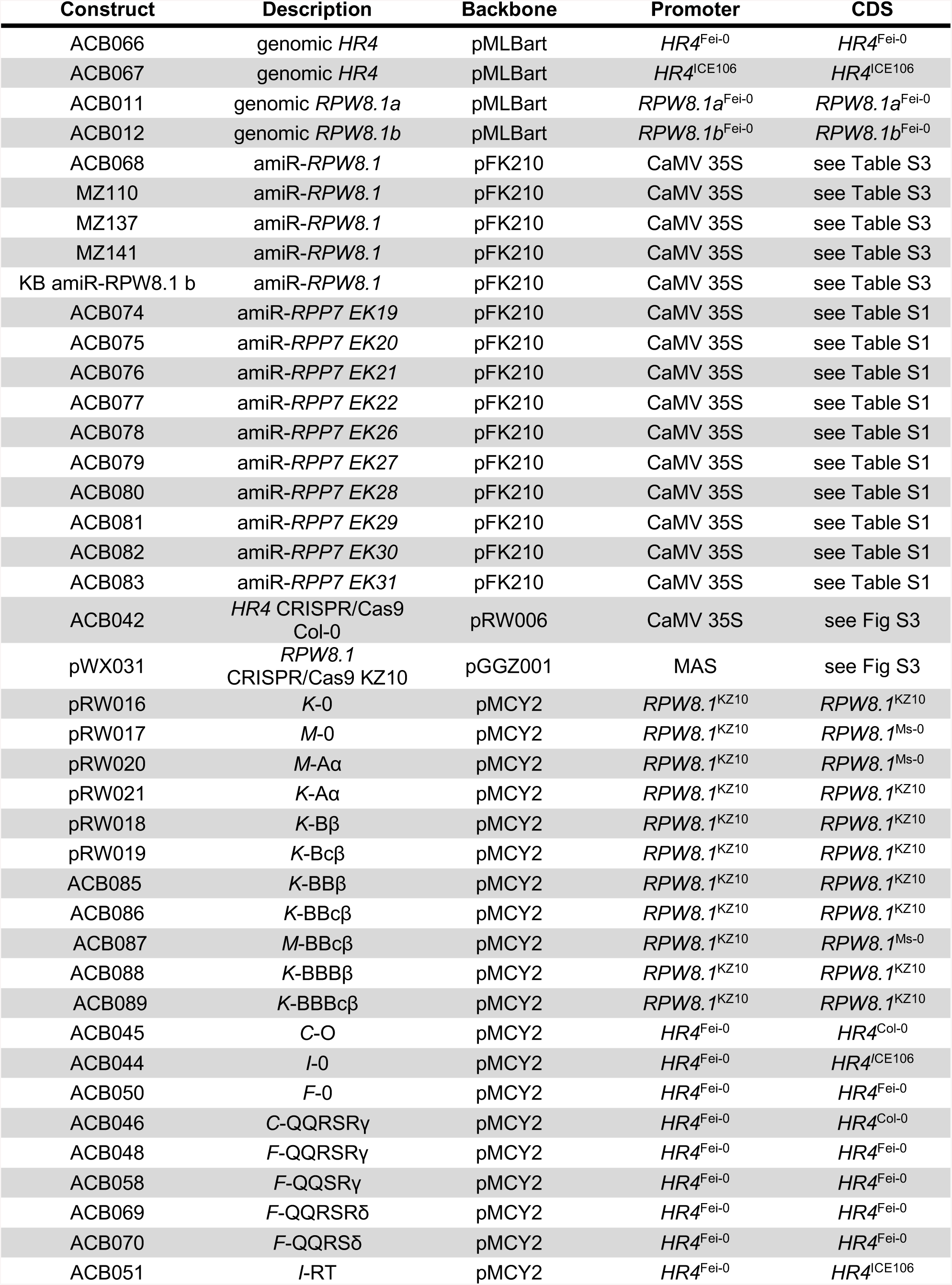
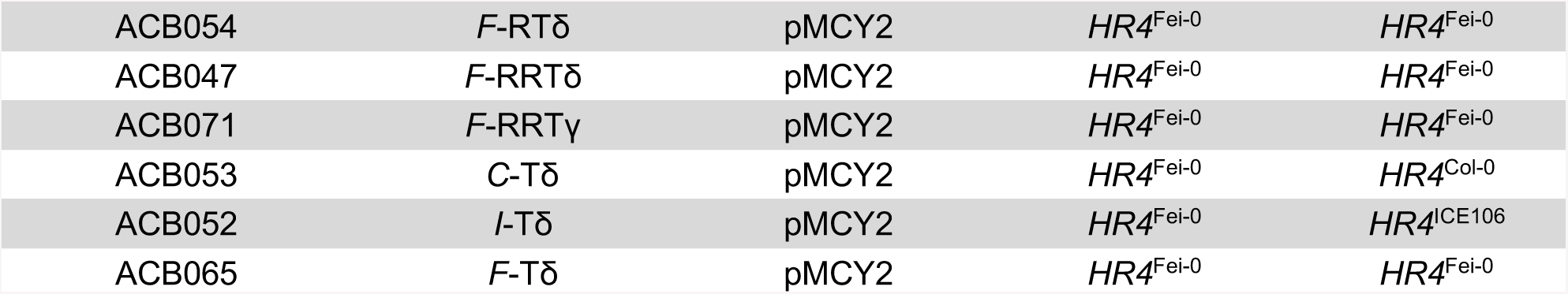
Constructs.

**Table S11.**
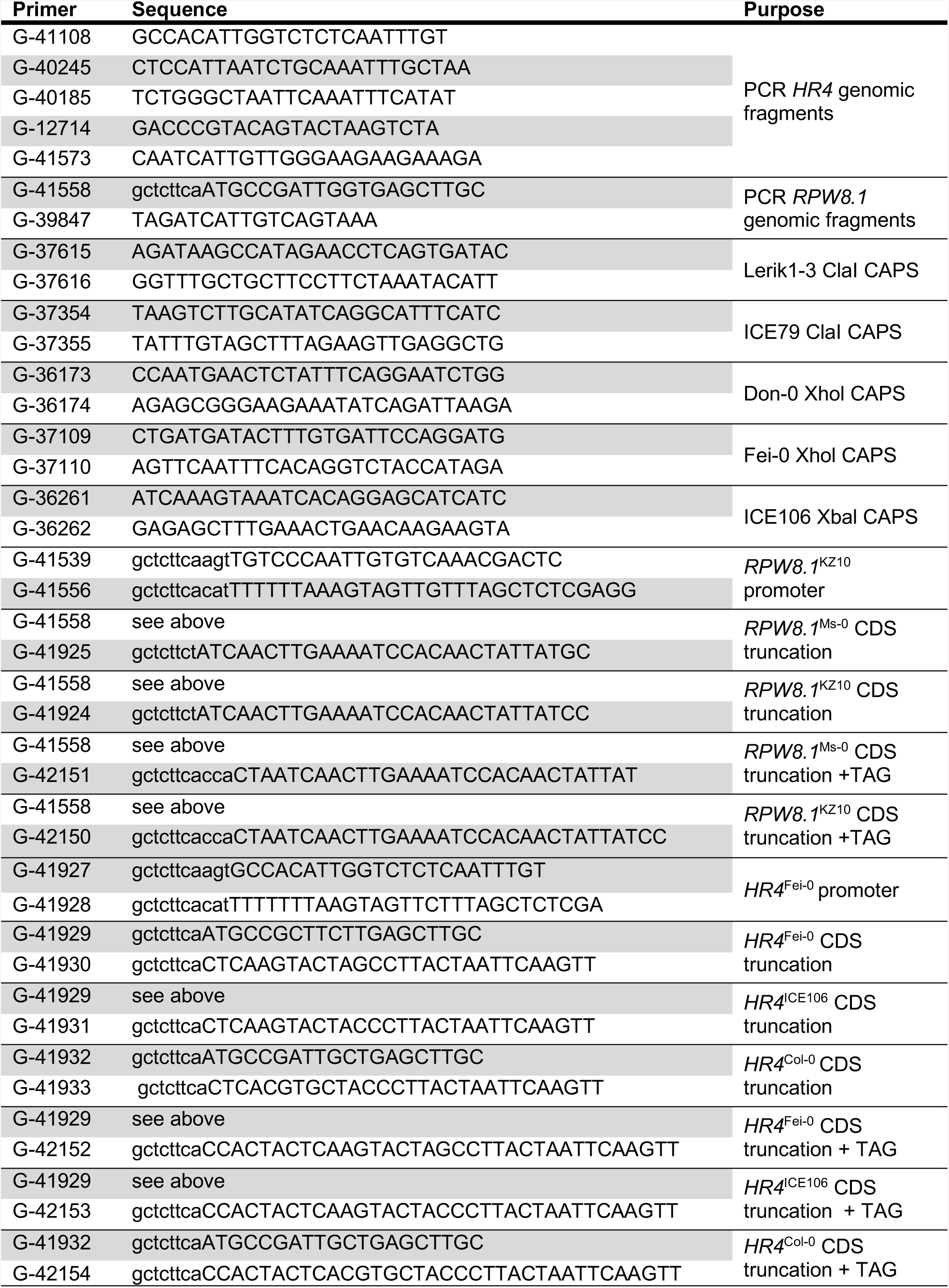
Oligonucleotides used for amplifying *RPW8.1/HR4* genomic fragments and swap constructs. Related to Fig 3 and Fig 5.

### Supplemental Experimental Procedures

#### *RPP7* phylogeny

The NB domain was predicted using SMART (http://smart.embl-heidelberg.de/). NB amino acid sequences were aligned using MUSCLE [1]. A maximum-likelihood tree was generated using the BLOSUM62 model in RaxML [2]. Topological robustness was assessed by bootstrapping 1,000 replicates.

#### RAPA phenotyping

Images were acquired daily in top view using two cameras per tray. Cameras were equipped with OmniVision OV5647 sensors with a resolution of 5 megapixels. Each camera was attached to a Raspberry Pi computer (Revision 1.2, Raspberry Pi Foundation, UK) [3]. Images of individual plants were extracted using a predefined mask for each plant. Segmentation of plant leaves and background was then performed by removing the background voxels then a GrabCut-based automatic postprocessing was applied [4]. Lastly, unsatisfactory segmentations were manually corrected. The leaf area of each plant was then calculated based on the segmented plant images.

#### Pathology

The *Hyaloperonospora arabidopsidis* isolate Hiks1 was maintained by weekly subculturing on susceptible Ws-0 *eds1-1* plants [5]. To assay resistance of susceptibility, 12- to 13-day old seedlings were inoculated with 5 x 10^4^ spores/ml. Sporangiophores were counted 5 days after infection.

#### Constructs and transgenic lines

Genomic fragments were PCR amplified, cloned into pGEM®-T Easy (Promega, Madison, WI, USA), and either directly transferred to binary vector pMLBart or Gateway vectors pJLBlue and pFK210. amiRNAs [6] against members of the *RPP7* and *RPW8*/*HR* clusters were designed using the WMD3 online tool (http://wmd3.weigelworld.org/), and placed under the CaMV 35S promoter in the binary vector pFK210 derived from pGreen [7]. amiRNA constructs were introduced into plants using *Agrobacterium*-mediated transformation [8]. T_1_ transformants were selected on BASTA, and crossed to incompatible accessions. For the chimeras, promoters and 5’ coding sequences were PCR amplified from genomic DNA, repeat and tail sequences were synthesized using Invitrogen’s GeneArt gene synthesis service, all were cloned into pBlueScript. The three parts, promoter, 5’ and 3’ coding sequences, were assembled using Greengate cloning [9] in the backbone vector pMCY2 [10]. Quality control was done by Sanger sequencing. Transgenic T_1_ plants were selected based on mCherry seed fluorescence. For CRISPR/Cas9 constructs, sgRNAs targeting *HR4* or *RPW8.1* were designed on the Chopchop website (http://chopchop.cbu.uib.no/), and assembled using a Greengate reaction into supervector pRW006 (pEF005-sgRNA-shuffle-in [11] Addgene plasmid #104441). mCherry positive T_2_ transformants were screened for CRISPR/Cas9-induced mutations by Illumina MiSeq based sequencing of barcoded 250-bp amplicons. Non-transgenic homozygous T_3_ lines were selected based on absence of fluorescence in seed coats.

